# Developmental assembly of multi-component polymer systems through interconnected gene networks *in vitro*

**DOI:** 10.1101/2024.03.14.585044

**Authors:** Daniela Sorrentino, Simona Ranallo, Francesco Ricci, Elisa Franco

## Abstract

Living cells regulate the dynamics of developmental events through interconnected signaling systems that activate and deactivate inert precursors. This suggests that similarly, synthetic biomaterials could be designed to develop over time by using chemical reaction networks to regulate the availability of assembling components. Here we demonstrate how the sequential activation or deactivation of distinct DNA building blocks can be modularly coordinated to form distinct populations of self-assembling polymers using a transcriptional signaling cascade of synthetic genes. Our building blocks are DNA tiles that polymerize into nanotubes, and whose assembly can be controlled by RNA molecules produced by synthetic genes that target the tile interaction domains. To achieve different RNA production rates, we use a strategy based on promoter “nicking” and strand displacement. By changing the way the genes are cascaded and the RNA levels, we demonstrate that we can obtain spatially and temporally different outcomes in nanotube assembly, including random DNA polymers, block polymers, and as well as distinct autonomous formation and dissolution of distinct polymer populations. Our work demonstrates a way to construct autonomous supramolecular materials whose properties depend on the timing of molecular instructions for self-assembly, and can be immediately extended to a variety of other nucleic acid circuits and assemblies.

## INTRODUCTION

A key feature of biomolecular materials is their ability to operate out of equilibrium and adapt to fluctuating environmental conditions.^1,2^ A classical example is given by cytoskeletal networks^3^, which are composed of filamentous assemblies that form and dissolve dynamically in response to endogenous and exogenous signals.^4,5^ Filament assembly organization is orchestrated by complex cellular signaling and regulatory networks that have evolved to interact synergistically, and determine when and where filaments will form.^6^

Learning from Nature, artificial biomolecular materials with the capacity to respond to stimuli have been built by rationally coupling components that self-assemble, and components that generate regulatory signals controlling the properties of self-assembling parts. In this context, approaches based on the use of nucleic acids to build both self-assembling elements and regulatory elements have been very productive.^7–9^ Due to their predictable non-covalent interactions that can be programmed through sequence design, DNA and RNA are ideal materials to build assemblies whose structural precision and complexity is approaching that of natural molecular machines.^10,11,12^ At the same time, the binding kinetics and equilibria of nucleic acid molecules can be prescribed by tuning the affinity of base-pairing domains,^13,14^ making it possible to build molecular systems operating like logic^15,16^ or dynamic circuits,^17^ and the presence of energy-dissipating enzymes or catalytic processes can further expanded the landscape of achievable dynamic behaviors.^14,18^ Naturally, nucleic acid assemblies and nucleic acid circuits can be seamlessly integrated: molecules produced by a nucleic acid circuit can be designed to hybridize with the domains of a nanostructure, thereby influencing its stability and capacity to assemble. This principle has fueled the development of DNA and RNA machines and supramolecular materials operating under the control of DNA and RNA networks.^19,20^ As major progress is being made toward building multi-component dissipative networks,^16,21^ and multi-component structures,^22,23^ questions are emerging about how to design network-structure interactions that are scalable and robust, and can achieve autonomous temporal behaviors.

Here, we propose a method to coordinate the assembly of distinct DNA components using modular synthetic genes to generate a transcriptional signaling cascade^24^. To achieve this, we have developed a platform of genes and DNA building blocks that can be modularly interconnected without crosstalk. Each gene of the cascade produces an RNA output carrying specific instructions to control a particular self-assembling component. The RNA output production rate can be fine-tuned through careful design of the gene. Further, the time at which the RNA output is released depends on the gene position in the cascade. Our building blocks are DNA tiles that polymerize into nanotubes,^25^ and whose assembly behavior can be controlled by RNA molecules targeting the tile interaction domains.^7,26^ By using the same functional components, genes and tiles, but changing how genes are cascaded as well as their RNA output production speed, we demonstrate that we can obtain nanotube assembly outcomes that differ in space and time, including random DNA polymers, block polymers, and as well as distinct autonomous formation and dissolution of distinct polymer populations.

We envision that our approach can be immediately extended to a variety of other nucleic acid circuits and assemblies.^18,19^ Beyond nanotechnology, our results illustrate how to develop autonomous supramolecular materials whose properties depend on the timing of biochemically-released assembly instructions, so that the same components can be routed toward a different fate depending on how regulatory signals are integrated.^27^

## RESULTS AND DISCUSSION

### Activation and inhibition of DNA building blocks

We consider double crossover (DX) DNA tiles composed of five distinct strands (Fig. 2a).^25,28,29^ Tiles interact via 5-nucleotide (nt) sticky ends leading to spontaneous self-assembly of micron-long nanotubes^28,30^ that can be visualized by labeling individual tiles with fluorophores.^25^ Building on previous work, we modified the tiles to be activated or inhibited by sequence-specific RNA strands that can be produced by synthetic genes.^7,31,32^ In our approach, one of the tile sticky ends is modified to include a 7-nt toehold (Fig. 2a, black domain) that serves as a binding domain for a single-stranded synthetic RNA inhibitor (Fig. 2a, light blue domain) complementary to both the toehold and sticky-end sequence (12 nt). The inhibitor not only prevents tile self-assembly, but it can also disassemble formed nanotubes due to the presence of the toehold which enables invasion of one of the tile sticky-ends (invasion of a single sticky-end is sufficient as tile assembly is a cooperative process). Therefore, addition of inhibitor results in nanotube disassembly within minutes (Fig. 2a).^7,33^ To activate the tiles and restore their ability to self-assemble, we employ an RNA activator designed to displace the inhibitor strand (Fig. 2a). The DNA sequences forming a tile are arbitrarily designed through computer programs,^34,35^ so one can generate an expandable set of sequence-distinct tile populations that can be individually controlled by their specific RNA activators and inhibitors.^7,26^

**Figure 1:**
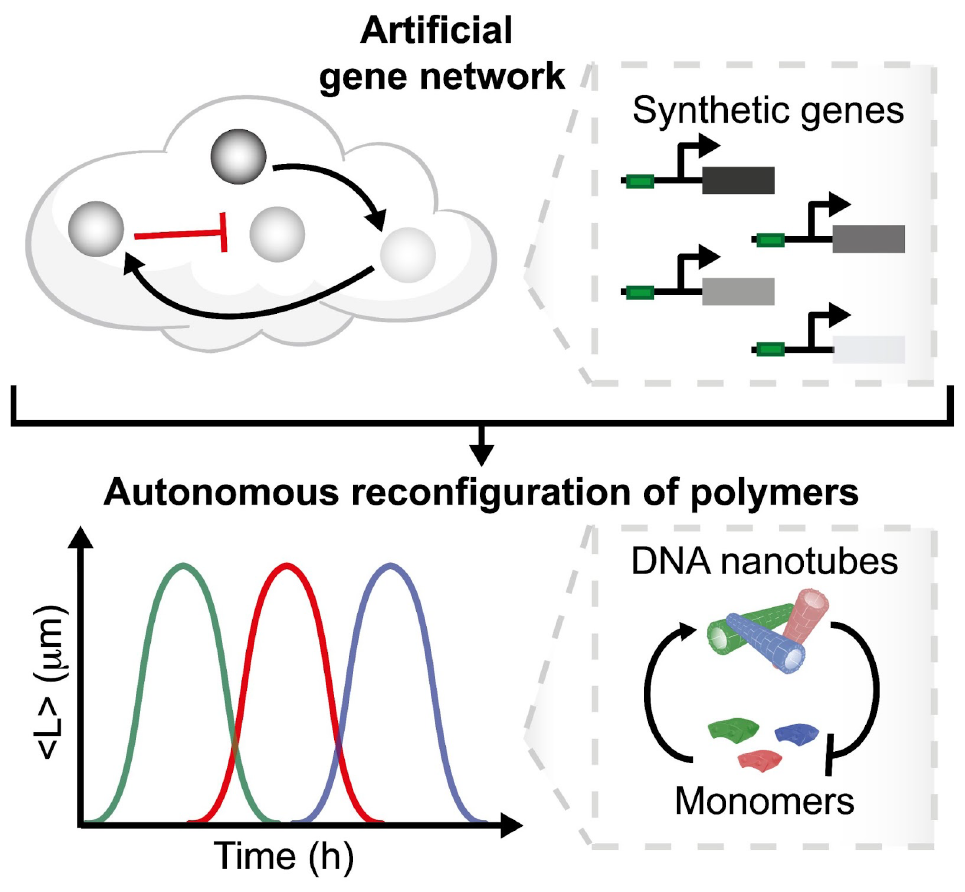
We integrate artificial gene networks to regulate the assembly of distinct building blocks, routing the system to different assembly states over time. We propose a scalable approach that integrates the principles of self-assembly and *in vitro* transcription to create a synthetic biopolymer system capable of programmed and autonomous reconfiguration. This is achieved through a suite of artificial genes and connectors that generate precise temporal instructions to activate or deactivate the assembly of distinct DNA-based monomers.

**Figure 2:**
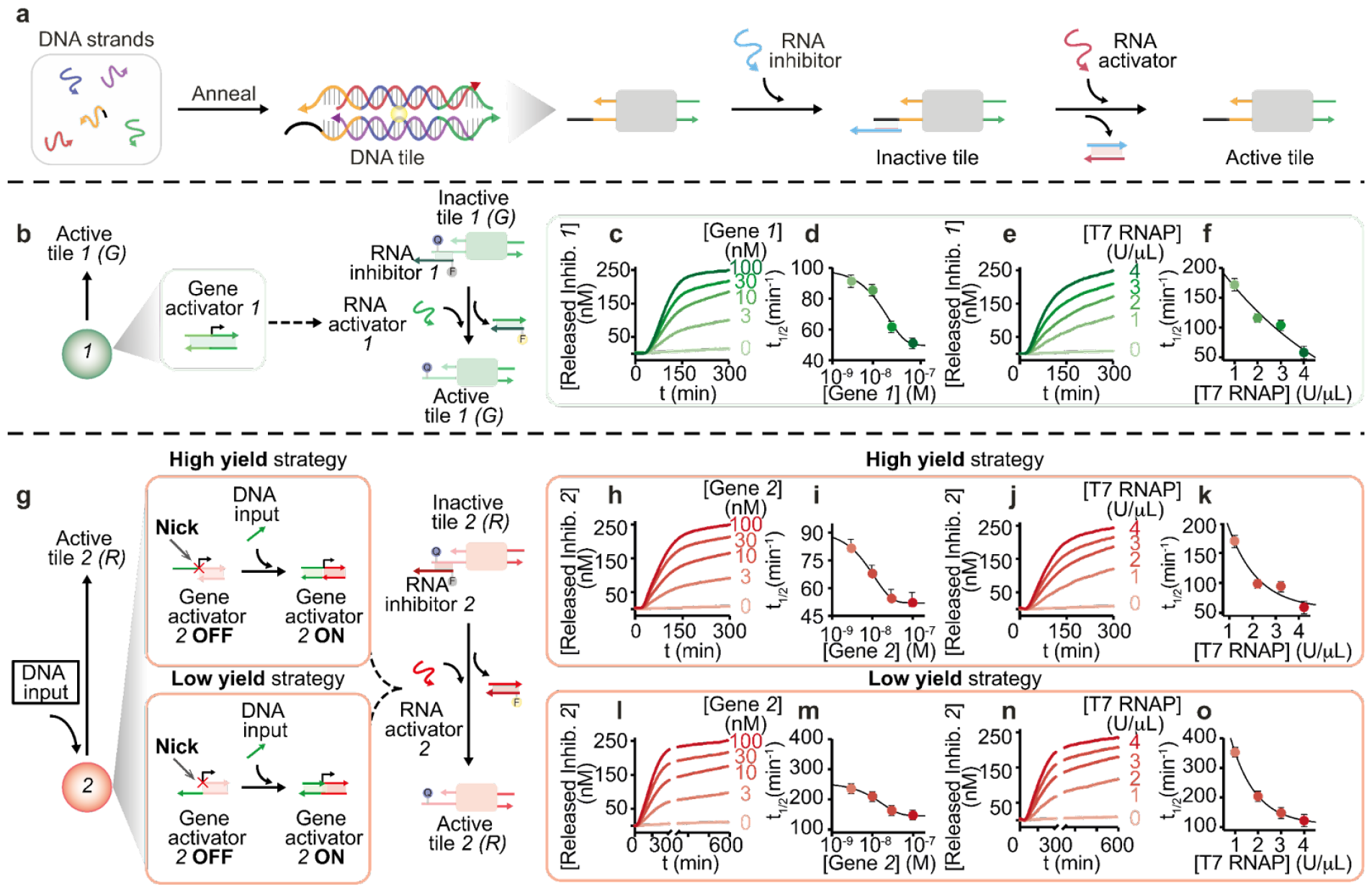
Optimization of RNA-induced activation of different DNA tiles. **a)** We consider DNA double crossover (DX) tile variants consisting of five unique DNA strands (here blue, red, yellow, green, and purple). The tiles self-assemble into nanotubes by hybridizing single-stranded complementary domains (i.e., sticky ends). Tiles were modified to contain a single-stranded overhang (toehold), shown as a black domain at the 5′-end of the yellow strand. Tiles can be represented schematically as molecular building blocks with complementary connectors. The presence of a toehold domain makes the adjacent sticky end accessible to strand invasion by an RNA inhibitor strand, thus inactivating the tiles and leading to nanotubes disassembly. By introducing an additional toehold domain into the RNA inhibitor strand, the tiles can be reactivated to reassemble using an RNA activator strand. In all sketches, the 3′-ends are marked with an arrow. **b)** Gene *1* (green filled circle) leads to transcription of RNA activator strand *1*, which activates tile *1*. **c)** Fluorescence kinetics at different concentrations of gene *1* (from 0 to 300 nM) showing displacement of RNA inhibitor *1* from inactive tile *1*. To easily follow the release of inhibitor from tiles after transcription of the RNA activator, the RNA inhibitor (1 μM) and one of the tile-forming strands are each labeled with a fluorophore/quencher pair (Cy3/BHQ1). **d)** Reaction half-activation time (t_1/2_) of inhibitor release from tile *1* as a function of gene *1* concentration. **e)** Fluorescence kinetics obtained at different concentrations of T7 RNAP (from 0 to 4 U/μL) added to a solution containing a fixed amount of tile *1* (250 nM), RNA inhibitor strand *1* (1 μM), and gene *1* (100 nM). **f)** Reaction half-activation time (t_1/2_) of RNA inhibitor release from tile as a function of T7 RNAP concentration. **g)** Gene *2* (red filled circle) promotes the transcription of RNA activator strand *2*, which in turn activates the corresponding tile *2*. Gene *2* is activated for transcription (ON) when a DNA input (green) binds and completes the T7 RNAP promoter sequence. Two different design strategies were used that altered the position of the nick on the promoter and resulted in different transcription yields. The high yield approach includes a nick on the template side, while the low yield approach includes a nick on the non-template side. **h)** Fluorescence kinetics at different concentrations (from 0 to 300 nM) of high yield synthetic gene *2* and equimolar amounts of DNA input (from 0 to 300 nM) to complete gene *2* show displacement of the RNA inhibitor from the inactive tiles by the presence of the transcribed RNA activator strand. To easily follow the release of the inhibitor from the tiles after transcription of the RNA activator, the RNA inhibitor (1 μM) and one of the tile-forming strands are each labeled with a fluorophore/quencher pair (Cy5/BHQ2). **i)** Reaction half-activation time (t_1/2_) of inhibitor release from DNA tile as a function of gene *2* concentration. **j)** Fluorescence kinetics obtained at different concentrations of T7 RNAP (from 0 to 4 U/μL) added to a solution containing a fixed amount of DNA tile *2* (250 nM), RNA inhibitor strand *2* (1 μM), gene *2* (100 nM), and DNA input (100 nM). **k)** Reaction half-activation time (t_1/2_) of RNA inhibitor *2* release from DNA tile as a function of T7 RNAP concentration. **l)** Fluorescence kinetics at different concentrations (from 0 to 300 nM) of low yield synthetic gene *2* and equimolar amounts of DNA input (from 0 to 300 nM) to complete gene *2*. To easily follow the release of the inhibitor from the tiles after transcription of the RNA activator, the RNA inhibitor (1 μM) and one of the tile-forming strands are each labeled with a fluorophore/quencher pair (Cy5/BHQ2). **m)** Reaction half-activation time (t_1/2_) of inhibitor release from DNA tile as a function of gene *2* concentration. **n)** Fluorescence kinetics obtained at different concentrations of T7 RNAP (from 0 to 4 U/μL) added to a solution containing a fixed amount of DNA tile *2* (250 nM), RNA inhibitor strand *2* (1 μM), gene *2* (100 nM), and DNA input (100 nM). **o)** Reaction half-activation time (t_1/2_) of inhibitor *2* release from DNA tile as a function of T7 RNAP concentration. Experiments shown in this Fig. were performed in 1X TXN buffer (5X contains: 200 mM Tris-HCl, 30 mM MgCl_2_, 50 mM DTT, 50 mM NaCl, and 10 mM spermidine), pH 8.0, 30 °C. Experimental values represent averages of three separate measurements and error bars reflect standard deviations.

### Synthetic genes can be designed to precisely tune the kinetics of tile activation

We developed synthetic genes producing RNA activators and inhibitors *in situ* that target a specific tile and control its active or inactive state with the desired kinetics. Each gene is a linear template that includes the T7 RNA polymerase (T7 RNAP) promoter site (Fig. 2b) and a downstream region encoding its RNA transcript sequence (RNA output).^36,37^ The speed of tile activation or inhibition can be modulated by changing the speed of RNA transcription. We illustrate this idea by measuring the kinetics of activation of DNA tile type *1* as a function of inhibitor, gene and T7 polymerase concentration (Fig. 2b) (RNA-based tile inhibition has been characterized in ^7^). The DNA tile is initially bound to its synthetic RNA inhibitor *1* and is activated by an RNA activator *1*. To assess the kinetics of activation, we labeled the RNA inhibitor strand with a fluorophore (Cy3) and the corresponding tile strand with a quencher (BHQ1). When the tiles are inactive, the strands labeled with the fluorophore and the quencher are in close proximity, resulting in a low fluorescence signal; fluorescence increases when the labeled inhibitor is displaced by transcribed RNA activator, which activates the tiles as shown in Fig. 2b. First, we checked in a control experiment that the addition of the synthetic RNA activator strand *1* to the inactive tiles causes their activation, as shown in Supplementary Information (SI Fig. 4). The kinetics of tile activation are captured by a simple ordinary differential equation model (Supplementary Note 1), using unfitted reaction rate parameters that are comparable with values found in the literature^7^ (SI Fig. 30). Next, given a fixed amount of tiles (250 nM) and RNA inhibitor (1 µM), we controlled the tile activation speed by varying the concentration of gene producing RNA activator (from 0 to 100 nM), obtaining a clear concentration-dependent response (Fig. 2c), with half-activation time that decreases with gene activator amount (t_1/2_= 50 ± 5 min at 100 nM gene *1* concentration) (Fig. 2d). After 24 hours, the inhibitor is fully released across all conditions (SI Fig. 6). Similarly, given a fixed amount of activator gene (100 nM), we can vary the T7 RNA polymerase level (from 0 to 4 U/μL) to modulate the speed of tile activation, achieving half-activation times between ∼50 and 180 min as shown in Fig. 2e,f. These experiments were also generally reproduced by a simple ODE model (SI Fig.s 31, 32), which suggests fast kinetic parameters for binding of RNAP and for tile activation.

Synthetic genes can be activated or deactivated through a promoter-displaced mechanism.^38,39^ In this approach (Fig. 2g), a synthetic gene can take one of two different states, OFF (inactive) or ON (active). The gene state depends on whether the promoter is double stranded or partially single stranded: one of the strands of the promoter site is nicked at -12, and if the upstream region is removed the gene is in an OFF state (negligible transcription). Transcription is switched ON if an ssDNA activator binds to and completes the promoter. When the gene is ON, we found that the transcription rate and yield depend on whether the nick is placed on the template strand (high yield) or on the non-template strand (low yield) (SI Fig.s 1-3). We take advantage of nick placement as a simple means to tune the kinetics of RNA transcription, and DNA tile activation, without altering the promoter sequence.

In Fig. 2g we demonstrate that a switchable synthetic gene can be used to obtain tunable tile activation kinetics; here we used a set of DNA tiles (red, *2*) that have the same sticky ends as tile *1* (see above) but a different toehold binding domain that allows recognition of a different synthetic RNA inhibitor strand *2*. Like done earlier, we labeled the RNA inhibitor *2* with a fluorophore (Cy5) and one of the tile strands with a quencher (BHQ2) to track the activation kinetics (Fig. 2g). In a control experiment, we verified that addition of RNA activator *2* successfully activates the tiles (SI Fig. 5). Next, we tested activation with switchable synthetic genes. When using the high yield gene *2* (nick on the template strand) we found that the speed of tile activation is comparable to the case in which we used constitutively active gene (Fig. 2b), both when we varied activator gene level and T7 RNAP level (DNA tiles *2* fixed at 250 nM and RNA inhibitor strand *2* at 1 μM) (Fig. 2h-k). When using the low yield gene *2* (nick on the non-template strand) we measured significantly slower tile activation kinetics, consistent with expectation given a much slower production of RNA activator. At high concentrations of gene *2* (100 nM) (Fig. 2l,m) and T7 RNAP (4 U/μL) (Fig. 2n,o), the tile half-activation time roughly doubles (t_1/2_ = 120 ± 20 min) when compared to the high yield design. All inhibitor is released after 24 hours across conditions (SI Fig. 7).

To summarize, we achieved fine control over tile activation kinetics by changing the transcription speed, and we engineered switchable genetic elements to further modulate activation.

### The outputs of a genetic cascade activate distinct tiles at specific times

Natural and artificial gene networks can autonomously generate a variety of dynamic behaviors. A gene cascade, in which one gene regulates the next in a chain, is a simple architecture to generate a temporal sequence of events synchronized with the activation of each gene. Here we build a simple cascade of two synthetic genes to sequentially control the activation of two DNA tile types. Building on the experiments we just described, we used a DNA “connector” complex to interconnect the genes *1* and *2* (Fig. 3a), which control tiles *1* and *2* respectively. Gene *1* is constitutively active, and it produces the RNA activator *1*. This RNA activator plays two important roles: it activates DNA tile *1*, and it simultaneously interacts with the connector complex to release a DNA molecule that activates the downstream gene *2* (Fig. 3a, SI Fig. 8). An advantage of this approach is that we can control the time it takes to activate tile *2* by modularly tuning various parameters of the genetic cascade.

**Figure 3:**
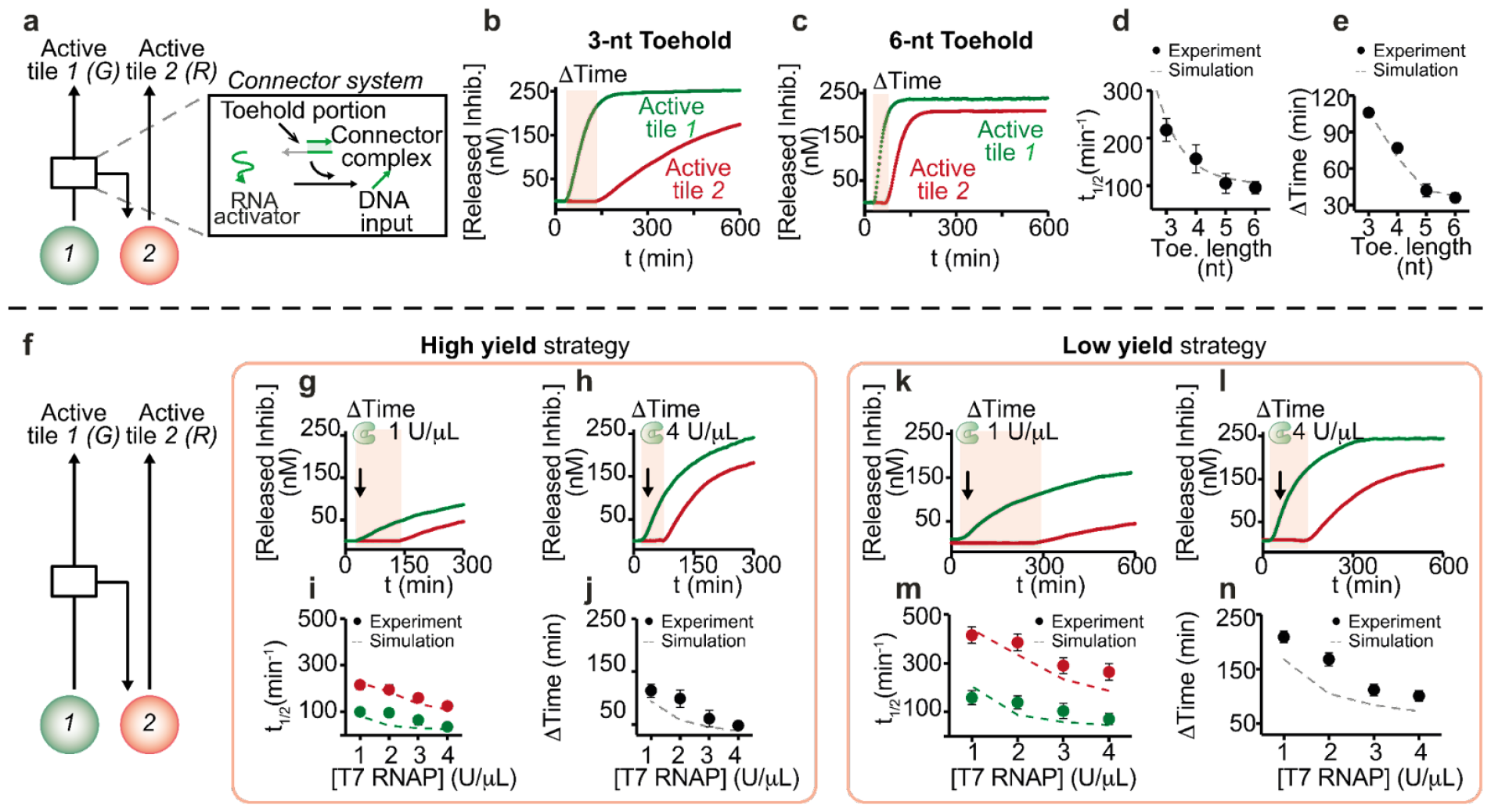
Sequential control of assembly of distinct tiles through a cascade of synthetic genes. **a)** Schematic representation of how two different genes (*1*, green circle and *2*, red circle) can be connected by a connector system. The second gene of the network is switched ON, when a DNA input is released from a connector complex. We designed one of the strands forming the connector complex to have the same 14-nt invading domain but differ in the length of the toehold region (from 3 to 6 nt). **b**,**c)** Fluorescence kinetics of the release of RNA inhibitor strands *1* and *2* from their corresponding tiles by the presence of transcribed RNA activator strands *1* and *2*, and connector complex at different toehold lengths (from 3 to 6 nt). To easily follow the release of the inhibitors from the tiles, the RNA inhibitor (1 μM) and one of the tile-forming strands were each labeled with a fluorophore/quencher pair (Cy3/BHQ1 and Cy5/BHQ2 for tile 1 and 2, respectively). **d)** Reaction half-activation time (t_1/2_) of inhibitor *2* released from DNA tiles *2* as a function of toehold length. **e)** Delay of inhibitor *2* release as a function of toehold length. **f)** The connector system allows sequential activation of two different tile types, each connected to a different gene (*1*, green, and *2*, red), occurring at different times. **g**,**h)** Fluorescence kinetics at different concentrations of T7 RNAP (from 0 to 4 U/μl) showing the displacement of RNA inhibitors *1* and *2* from the corresponding tiles by the transcribed RNA activators, using a fixed concentration of high yield gene 2 (100 nM), connector complex (300 nM), and DNA input (300 nM). **i)** Reaction half-activation time (t_1/2_) of inhibitors *1* (green) and *2* (red) released from DNA tiles as a function of T7 RNAP concentration. **j)** The delay time at which tile *2* activation begins can be tuned by varying the concentration of T7 RNAP. **k**,**l)** Fluorescence kinetics at different concentrations of T7 RNAP (from 0 to 4 U/μl) showing the displacement of RNA inhibitors *1* and *2* from the corresponding inactive tiles by the presence of the transcribed RNA activators, using a fixed concentration of low yield gene *2* (100 nM) and connector complex (300 nM), and DNA input (300 nM). **m)** Released inhibitor from tiles *1* and *2* as a function of T7 RNAP concentration at 600 min reaction. **n)** Time-dependent response of tile activation at different concentrations of T7 RNAP. Experiments shown in this Fig. were performed in 1X TXN buffer (5X contains: 200 mM Tris-HCl, 30 mM MgCl_2_, 50 mM DTT, 50 mM NaCl, and 10 mM spermidine), pH 8.0, 30 °C. Experimental values represent averages of three separate measurements and error bars reflect standard deviations. In panels **d, e, i, j, m**, and **n** dots represent experimental values, while dashed lines represent fits obtained with the mathematical model.

First, we adapted the connector design to tune the activation speed and the activation onset delay (ΔT) of tile *2*. We evaluated connector complexes that have the same 14-nt invading domain but differ in the length (3-6 nt) of the toehold that initiates the release of the gene DNA activator (Fig. 3a). We used conditions consistent with previous experiments (250 nM each tile *1* and *2*, 1 μM RNA inhibitors *1* and *2*, and 100 nM high yield synthetic genes *1* and *2*). To easily follow the kinetics of tile activation, we labeled the RNA inhibitors with a fluorophore (Cy3 and Cy5 for inhibitors *1* and *2*, respectively) and the tiles with their corresponding quenchers (BHQ1 and BHQ2). The speed of tile activation and the delay ΔT strongly depend on the toehold length (Fig. 3b,c and SI Fig. 9). Indeed, the half-activation time (t_1/2_) for tile *2* increases from 95 to 215 minutes when the length of the toehold domain is reduced from 6 to 3 nt (Fig. 3d,e and SI Fig. 9). We verified that the DNA activator for gene *2* is displaced much faster from the connector complex by RNA output *1* when compared to the displacement of inhibitor from tile *2* (tile activation requires the additional transcription of RNA *2*) (SI Fig.s 10,11). We expanded our kinetic model of tile activation and activator production to capture this interconnected system (Supplementary Note 1), and were able to reproduce the measured t_1/2_ and ΔT as the connector strand displacement speed changes with the toehold length (Fig. 3 d,e dashed lines). For the next experiments we designed connectors with the fastest strand displacement, *i*.*e*. 6 nt toehold.

Next, we modulated the RNA transcription speed of gene *2* to influence the activation kinetics of tile *2*, taking advantage of the high and low yield gene designs characterized earlier (Fig. 3g). Like before, we used fluorescently-labeled RNA inhibitors *1* and *2* to track their release from tiles, and we monitored their level over time under variable T7 RNAP concentration (from 1 to 4 U/μL). When using high yield genes, the release of RNA inhibitors (*1*, green and *2*, red) is complete after about 180 min at high T7 RNAP level (1U/µL), while the release is not completed at low T7 RNAP concentration (1 U/μL) even after 300 min (Fig. 3h). Furthermore, by using different T7 RNAP concentrations we can also control the delayΔT in the onset of RNA production (Fig. 3i,j). When changing the T7 RNAP concentration (1 to 4 U/µL) we observed ΔT values ranging from 120 to 50 minutes (Fig. 3j, SI Fig. 12). When using low yield genes, we observe a more significant ΔT of 240 minutes at low T7 RNAP concentration (1 U/μL), and even after 600 minutes we do not achieve a complete release of RNA inhibitors (Fig. 3k). As before, using a higher T7 RNAP concentration improves the release speed and reduces the delay ΔT (Fig. 3k-n, SI Fig. 13). We finally tested tile activation speed when the concentration of one of the genes is kept constant, and that of the other gene varies. As expected, when the concentration of gene *2* (downstream gene in the cascade) is held constant (100 nM), as we increase the concentration of gene *1* (from 3 to 100 nM) we obtain a faster release of the RNA inhibitors while the delay ΔT decreases (SI Fig.s 14-16). Naturally, when the concentration of gene *1* is constant, the release of RNA inhibitor *1* remains unchanged and does not depend on the concentration of the downstream gene *2*; in contrast, the release of RNA inhibitor *2* depends on the concentration of gene *2*, with more inhibitor being released at higher concentrations (SI Fig.s 17–23). Also in this case, the measured t_1/2_ and ΔT are reproduced by an unfitted kinetic model in which the parameters for tile activation and transcription are consistent across all simulations^7,40^; the low yield strategy is captured by assuming slower binding and faster unbinding of RNAP to the gene, when compared to the high yield case. The agreement of the simulations and experimental data confirms that the kinetics of the components involved are generally predictable and modular.

### Timing assembly and polymer organization through different genetic cascades

We showed how distinct tile types can be activated at different times by modulating the features of an autonomous two-gene cascade. Next, we examine how this fine temporal control over tile activation impacts the dynamics of tile assembly into nanotubes, as well as the properties of the nanotubes. Using the tile designs introduced earlier (*1* and *2*), we characterized the kinetics of nanotube growth in the presence of the two-gene cascade. In these experiments each tile carries a different molecular “load”, as it is decorated with a distinct fluorophore (tile 1 with Cy3, nanotubes colorized in green; and tile 2 with Cy5, nanotubes colorized in red) which makes it possible to track assembly via fluorescence microscopy. As in prior experiments, each tile (250 nM) is initially bound to its RNA inhibitor (1 µM), and activated by their transcribed RNA activator.

We first verified that genes *1* (constitutively active) and *2* (switchable) individually trigger the assembly of micron-long DNA nanotubes. Fig. 4a and b show that gene *1* correctly induces assembly of green nanotubes from tile *1*, by producing RNA 1 that activates tile *1*. These nanotubes present a mean length of 3–4 µm after about 8 hours, while their density (number of nanotubes per 100 μm^2^) decreases over time due to joining events^30^ (Fig. 4c,d). It is noteworthy that the temporal evolution of the growth of the nanotubes influences the count (i.e. the number of structures 100 μm^2^). As the average length of the nanotubes increases, their number decreases proportionally to the transcription time (Fig. 4c,d). Next, we verified that gene *2* correctly induces assembly of tile *2* into red nanotubes, in both the high and low yield variants. As expected, the high yield variant of gene *2* induces nanotube growth significantly faster than the low yield variant. Nanotubes reach their plateau length within 8 hours when controlled by the high yield gene (Fig. 4f-h). It takes more than 12 hours to reach a comparable mean length under the control of the low yield gene (Fig. 4i-k), which introduces a noticeable time delay before nanotubes are visible, due to the longer time it takes to build up enough RNA to activate tiles above their nucleation threshold.

**Figure 4:**
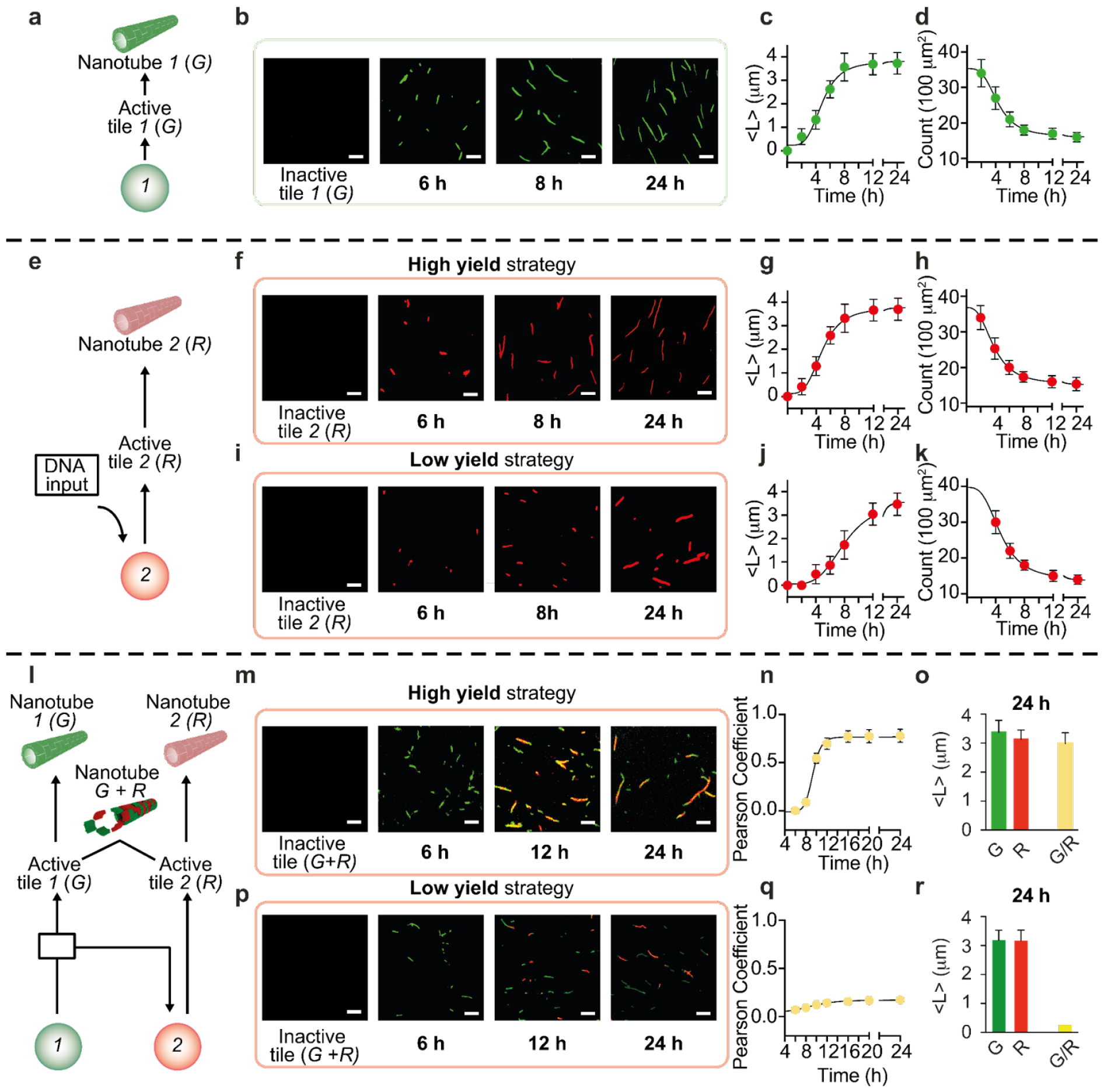
Controlling the growth and composition of DNA nanotubes through individual and interconnected genes. **a)** Gene *1* (green circle) activates the corresponding tile *1* to self-assemble into green DNA nanotubes by producing the specific RNA activator *1*. **b)** Fluorescence microscopy images show self-assembly of green (G) nanotubes after production of RNA activator in the presence of T7 RNAP (4 U/μl). No nanotubes are observed in the presence of the specific RNA inhibitor. Nanotubes labeled with Cy3 dye were imaged at a concentration of 50 nM. **c)** Kinetic traces of nanotube length measured from fluorescence microscopy images; <L>indicates mean nanotube length. **d)** Count (number of structures per 100 μm^2^) of assembled green nanotubes measured from fluorescence microscopy images. **e)** Gene 2 (red circle), when completed with an external DNA input, leads to the activation of tile R, which resembles red DNA nanotubes. **f)** Fluorescence microscopy images showing self-assembly of red (R) nanotubes after production of RNA activator using a high yield gene in the presence of T7 RNAP (4 U/μl). No nanotubes are observed in the presence of the specific RNA inhibitor. Nanotubes labeled with Cy5 dye were imaged at a concentration of 50 nM. **g)** Kinetic traces of nanotube length measured from fluorescence microscopy images. <L>indicates the mean nanotube length of nanotubes. **h)** Count (number of structures per 100 μm^2^) of assembled red nanotubes measured from fluorescence microscopy images. **i)** Fluorescence microscopy images showing self-assembly of red (R) nanotubes after production of RNA activator using a low yield gene in the presence of T7 RNAP (4 U/μl). No nanotubes are observed in the presence of the specific RNA inhibitor. Nanotubes labeled with Cy5 dye were imaged at a concentration of 50 nM. **j)** Kinetic traces of nanotube length measured from fluorescence microscopy images. <L>indicates the mean nanotube length of nanotubes. **k)** Count (number of structures per 100 μm^2^) of assembled red nanotubes measured from fluorescence microscopy images. **l)** Interconnected genes *1* (green circle) and 2 (red circle) allow sequential activation of two different tiles (G and R) to drive the assembly of two different nanotube populations. **m)** Fluorescence microscopy images show self-assembly of green (G) and red (R) random nanotubes after production of the two RNA activators from their respective genes (gene 2 is a high yield) in the presence of T7 RNAP (4 U/μl). No nanotubes are observed in the presence of the specific RNA inhibitors. Nanotubes labeled with Cy3 (G) and Cy5 (R) dyes were imaged at a concentration of 50 nM. **n)** Pearson coefficient values of reassembled random G/R nanotubes. **o)** Histograms of nanotube length (<L>/μM) measured from fluorescence microscopy images for each channel (green, G, red, R, and merged, G/R) after the 24-hour reaction. **p)** Fluorescence microscopy images showing self-assembly of green (G) and red (R) nanotubes after production of the two RNA activators from their respective genes (gene 2 is a low yield) in the presence of T7 RNAP (4 U/μl). No nanotubes are observed in the presence of the specific RNA inhibitors. Nanotubes labeled with Cy3 (G) and Cy5 (R) dyes were imaged at a concentration of 50 nM. **q)** Pearson’s coefficient values of reassembled G/R nanotubes. **r)** Histograms of nanotube length (<L>/μM) measured from fluorescence microscopy images for each channel (green, G, red, R, and merged, G/R) after the 24-hour reaction. Experiments shown in this Fig. were performed in 1X TXN buffer (5X contains: 200 mM Tris-HCl, 30 mM MgCl_2_, 50 mM DTT, 50 mM NaCl, and 10 mM spermidine), pH 8.0, 30 °C. [Tile G] = [Tile R] = 250 nM; [RNA inhibitors] = 1 μM; [gene 1] = [gene 2] = 100 nM; [connector complex]= 300 nM. Experimental values were calculated via triplicate experiments, and error bars reflect standard deviations. Scale bars for all microscope images, 2.5 µm.

We finally controlled the self-assembly of the two tile types simultaneously, under the control of our interconnected two-gene cascade (Fig. 4l). We used the constitutively active gene *1* to activate either the high yield or low yield version of gene 2 (100 nM each gene), using our connector complex (300 nM). As the cascaded genes produce their RNA activators sequentially, tiles *1* and *2* also self-assemble into nanotubes sequentially. Because tiles *1* and *2* have identical sticky ends and can interact, by tuning the speed of the cascade we can control the level of free tiles that are active at a particular point in time. If tiles *1* and *2* are activated nearly simultaneously, nanotubes should assemble from a mix of tiles yielding random green-red polymers. If tile *2* is activated much later than tile *1*, green nanotubes should form first depleting free tile *1*, making it possible for tile *2* to assemble into red nanotubes: in this case, we should obtain block green or red polymers. We built a fast cascade using the high yield version of gene *2*: this results in rapid activation of tiles *2*, which combine with unpolymerized tile *1* and yield random polymers that appear yellow in the merged channels (Fig. 4m-o). A slow cascade including the low yield version of gene *2* results in delayed and slower activation of tile *2*, which exceeds the threshold for nucleation when tiles of type *1* are completely polymerized into green nanotubes. Thus, tile *2* produces red nanotubes which begin to be visible around 12 h (Fig. 4p-r). We characterized quantitatively the nanotube composition under the control of the fast and slow cascade by tracking the statistical properties of epifluorescence microscopy images in terms of the spatial localization of individual tiles within nanotubes. We calculated the Pearson coefficient (PC), which measures the strength of the linear relationship between the fluorescence intensity values of the green and red regions on the nanotubes. PC Values close to 0 indicate low colocalization of the fluorophores on the structures, while PC values around 1 indicate high colocalization. With the fast gene cascade yielding randomly distributed nanotubes, we measured a Pearson coefficient PC = 0.80 ± 0.16, which confirms colocalization (Fig. 4n,o). For the slow gene cascade, we obtained a PC of 0.20 ± 0.02, which indicates very limited colocalization of the tiles (Fig. 4q,r).

By combining fast and slow genetic components in a cascade we can thus program the timing as well as the properties of self-assembled structures, and this approach can be scaled to a larger number of genes and assembling components, as we will show in the next sections.

Building on the lessons learned from the two-gene cascade, we demonstrate a series of temporal programs to assemble diverse types of nanotubes through a three-gene cascade shown in Fig. 5a. Again, the genes are cascaded through connector complexes that bind to the RNA activator produced by the gene upstream, and release DNA activator for the gene downstream. Like before, each gene activates specifically only one type of tile through a unique activator (gene *1* activates tile *1*, colored in green; gene *2* activates tile *2*, colored in red, and gene *3* activates tile *3*, shown in blue), but tiles share their sticky ends and can interact. For this reason, like in the two gene cascade, the timing of activation of each tile type influences the random or block polymer organization of nanotubes.

**Figure 5:**
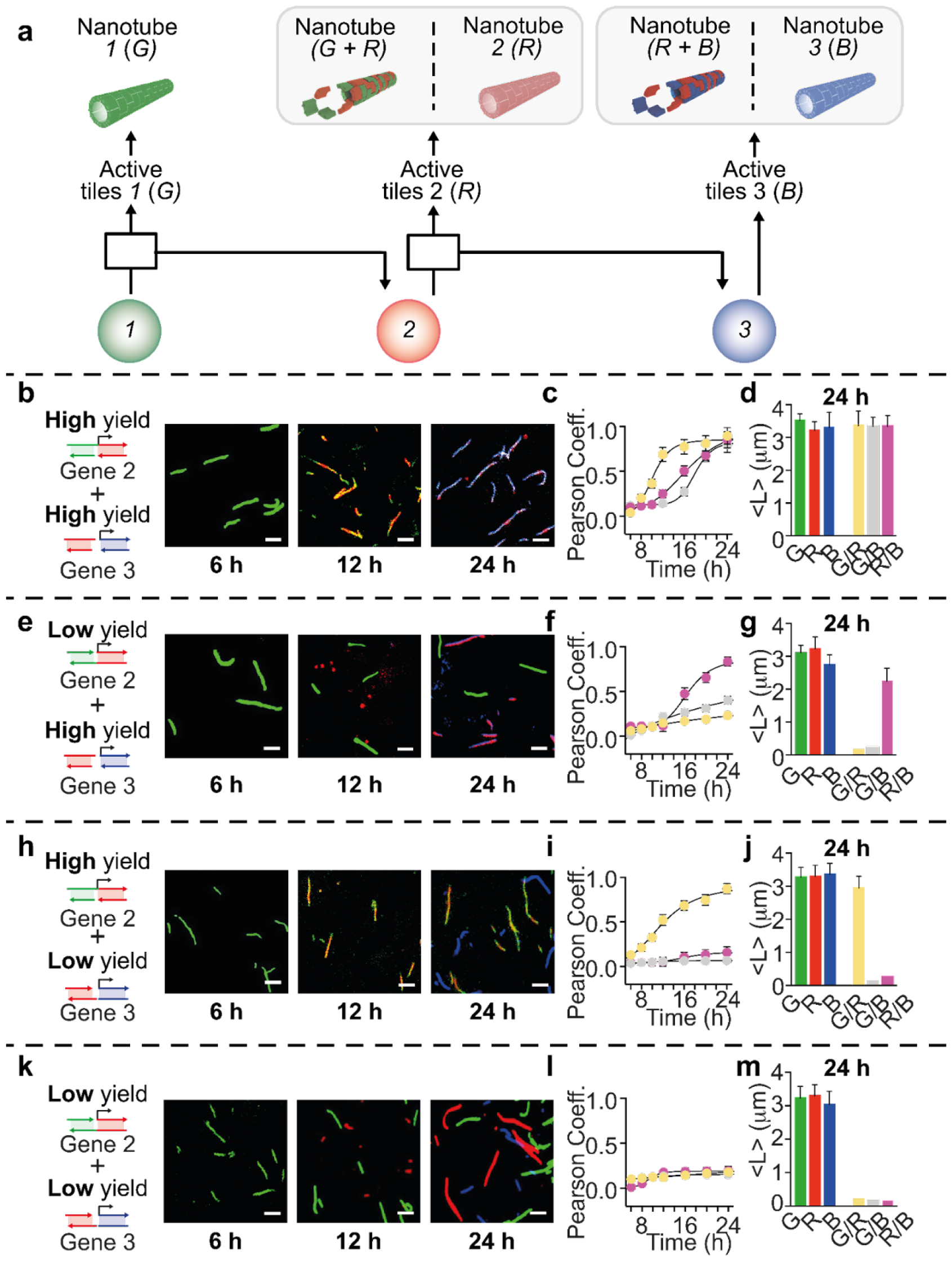
Development of different nanotube populations by tuning the production kinetics of distinct RNA activators. **a)** Three interconnected genes are activated by their respective upstream nodes at a specific time point, resulting in the formation of distinct nanotube populations. **b)** Fluorescence microscopy images showing self-assembly of green (G), red (R), and blue (B) nanotubes after production of the corresponding RNA activators in the presence of T7 RNAP (4 U/μl) using a combination of both high yield genes 2 and 3. **c)** Pearson coefficient of reassembled G/R/B nanotubes at different reaction times after addition of T7 RNAP (4 U/μl). **d)** Histograms of nanotube length (<L>/μM) measured from fluorescence microscopy images for each channel (green, G, red, R, blue, B and merged, G/R, G/B and R/B) after the 24-hour reaction. **e)** Fluorescence microscopy images showing self-assembly of green (G), red (R), and blue (B) nanotubes after production of their RNA activators in the presence of T7 RNAP (4 U/μl) using a combination of low yield gene 2 and high yield gene 3. **f)** Pearson coefficient of reassembled G/R/B nanotubes at different times after addition of T7 RNAP (4 U/μl). **g)** Histograms of nanotube length (<L>/μM) measured from fluorescence microscopy images for each channel (green, G, red, R, blue, B and merged, G/R, G/B and R/B) after the 24-hour reaction. **h)** Fluorescence microscopy images showing self-assembly of green (G), red (R), and blue (B) nanotubes after production of their RNA activators in the presence of T7 RNAP (4 U/μl) using a combination of high yield gene 2 and low yield gene *3*. **i)** Pearson coefficient of reassembled G/R/B nanotubes at different times after addition of T7 RNAP (4 U/μl). **j)** Histograms of nanotube length (<L>/μM) measured from fluorescence microscopy images for each channel (green, G, red, R, blue, B and merged, G/R, G/B and R/B) after the 24-hour reaction. **k)** Fluorescence microscopy images showing self-assembly of green (G), red (R), and blue (B) nanotubes after production of their RNA activators in the presence of T7 RNAP (4 U/μl) using a combination of both low yield genes 2 and *3*. **l)** Pearson coefficient of reassembled G/R/B nanotubes at different times after addition of T7 RNAP (4 U/μl). **m)** Histograms of nanotube length (<L>/μM) measured from fluorescence microscopy images for each channel (green, G, red, R, blue, B and merged, G/R, G/B and R/B) after the 24-hour reaction. Experiments shown in this Fig. were performed in 1X TXN buffer (5X contains: 200 mM Tris-HCl, 30 mM MgCl_2_, 50 mM DTT, 50 mM NaCl, and 10 mM spermidine), pH 8.0, 30 °C. [Tile G] = [Tile R] = [Tile B]= 250 nM; [RNA inhibitors] = 1 μM; [gene 1] = [gene 2] = [gene 3] 100 nM; [connector complexes]= 300 nM. Experimental values were calculated via triplicate experiments, and error bars reflect standard deviations. Scale bars for all microscope images, 2.5 µm. In **b, e, h**, and **k**, nanotubes labeled with Cy3 (G), Cy5 (R), and FAM (B) were imaged at a concentration of 50 nM. In **c, f, i**, and **l**, the G/R channel gives a yellow color, G/B gives a gray color, and R/B gives a pink color.

We demonstrate four different nanotube organization patterns by using different combinations of high-yield and low-yield genes that change the speed of activation of tiles in the cascade. Using high-yield variants of both genes *2* and *3*, we obtained nanotubes with a random distribution of the three different tiles. This is evident from the overlaps of the green, red and blue channels and is also confirmed by the calculated PC over time (Fig. 5b,c). The average length of the structures measured from the merged channels (G/R, G/B and R/B) is similar to the length measured from the individual channels, with values within the standard deviation (Fig. 5d). When a low-yield version of gene *2* is used, we observe limited co-localization of tiles *1* and *2* (green/red). However when combined with a high-yield version of gene *3*, the activation of tile *3* is nearly as fast as that of tile *2*, and we obtain random red/blue nanotubes (Fig. 5e-g). As expected, the combination of high-yield gene *2* and of low-yield gene *3* results in random green and red nanotubes (tiles *1* and *2*) but blue block polymers (tile *3*) (Fig. 5h-i). Finally, when using low-yield variants of both genes 2 and 3, only block polymers are formed since each tile type is activated slowly and is completely polymerized before the next tile in the chain becomes active (Fig. 5k-m). These experiments collectively illustrate how we can autonomously route the assembly of a material, here exemplified by DNA tiles loaded with different fluorogenic molecules, by taking advantage of interconnected gene networks and their programmable temporal responses. This approach could be scaled up to include more complex gene networks that could deliver a variety of instructions, beyond activation of the self-assembling monomers, as we show in the next section.

### Programming temporal waves of assembly of distinct nanotube populations

Temporal sequences of assembly and disassembly of biomolecular scaffolds enables many biological functions like cell growth, motility, and development. Here we take advantage of our suite of synthetic genes to control not only the appearance, but also the dissolution of multiple types of DNA nanotubes in an autonomous manner. We focus on demonstrating the transient, sequential assembly of three types of tiles each carrying a different fluorescent cargo. We engineered six cascaded genes, three of which produce RNA activators that enable tile assembly into nanotubes (genes 1-3), and the other three (genes 4-6) produce RNA inhibitors (Fig. 6a).

**Figure 6:**
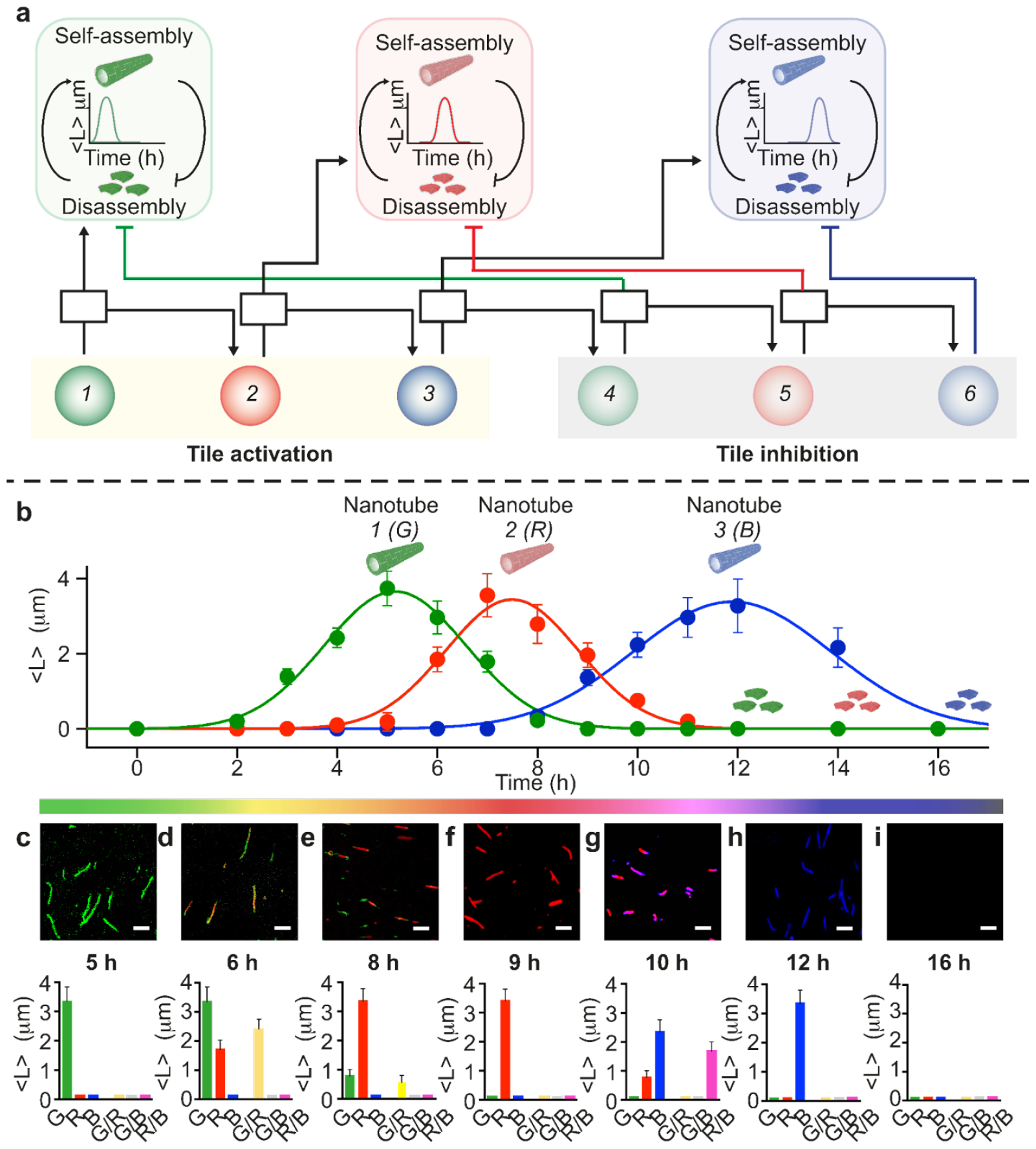
Autonomous, transient temporal waves of distinct nanotube populations driven by a genes cascade. **a)** We designed six interconnected genes (filled colored circles) that can control the self-assembly (the first three) or disassembly (the second three) of DNA nanotubes by transcribing specific RNA activators or inhibitors. **b)** When the specific RNA activator displaces the inhibitor from the corresponding tile, the activation process becomes dominant and promotes nanotube regrowth. In contrast, the production of RNA inhibitors leads to their gradual degradation. Kinetic traces show the autonomous assembly and disassembly of G, R, and B nanotubes (mean length, <L>) at different reaction time points. **c-i)** Fluorescence microscopy images showing that the mean nanotube length increases when growth is triggered by transcription of RNA activators and decreases when transcription of inhibitors begins. Histograms of nanotube length (<L>/μM) measured from fluorescence microscopy images for each channel (green, G, red, R, blue, B and merged, G/R, G/B and R/B). Experiments shown in this Fig. were performed in 1X TXN buffer (5X contains: 200 mM Tris-HCl, 30 mM MgCl2, 50 mM DTT, 50 mM NaCl, and 10 mM spermidine), pH 8.0, 30 °C; [Tile G] = [Tile R] = [Tile B]= 250 nM; [RNA inhibitors] = 1 μM; [gene 1] = [gene 2] = [gene 3] = 100 nM; [gene 4] = [gene 5] = [gene 6]= 3 nM; [connector complexes]= 300 nM; Scale bars for all microscope images, 2.5 µm. Nanotubes labeled with Cy3 (G), Cy5 (R), and FAM (B) were imaged at a concentration of 50 nM. In **b**-**i** the mean and s.d. of nanotube length are calculated over triplicate experiments. The G/R channel gives a yellow color, G/B a gray color, and R/B a pink color.

We programmed our cascade to produce RNA inhibitors with a significant delay when compared to the activation steps. This is due to the fact that the disassembly of nanotubes occurs much faster than their assembly upon tile activation ^7^. We delayed the inhibition genes by slowing down the response of the connector complexes. In a control experiment we verified that a reduced toehold length in the connector complex leads to an increase in the time delay for tile disassembly, with shorter toeholds leading to longer delays (SI Fig.s 24-26). This was verified by testing connector complexes with toeholds between 3 and 6 nt to release DNA activator for a set of new genes (numbered 4, 5 and 6) designed to produce RNA inhibitor for tiles 1, 2 and 3. We selected a toehold length of 3 nt for all connectors activating genes 4, 5 and 6 that produce RNA inhibitors (SI Fig.s 27-29).

Our six-gene cascade achieves sequential steps of assembly and disassembly of three distinct types of tiles. This autonomous succession of events was achieved by combining in one pot tiles 1, 2, and 3 (250 nM each) in a solution along with their corresponding RNA inhibitors (1 μM each), the complete set of connectors (300 nM each), and high yield gene variants (100 nM each). After addition of T7 RNAP, we observe rapid growth of green nanotubes resulting from the assembly of tile 1 (Fig. 6b,c). As gene 2 is activated, red nanotubes and random green/red nanotubes appear (Fig. 6d). Because the reactions involving connector complexes and RNA transcription proceed faster than self-assembly, we begin to observe disassembly of green nanotubes as gene 4 becomes active, before we can observe blue nanotubes (activated by gene 3) form (Fig. 6e,f). Blue nanotubes appear about 10 hours after the start of the reaction, and random red/blue polymers form (Fig. 6g). As gene 5 becomes active and produces inhibitor for tile 2, the red nanotubes begin to disassemble. Next, gene 6 becomes active and induces disassembly of blue nanotubes. After 16 hours the process is complete and all nanotubes have been dissolved (Fig. 6i).

## CONCLUSION

We have demonstrated a platform of modular components to build DNA-based materials whose properties evolve according to tunable temporal programs. Taking inspiration from how cells regulate sequential developmental events,^41^ we have used minimal artificial genetic cascades to produce RNA outputs that sequentially control activation or deactivation of distinct self-assembling DNA tiles. These RNA signals make it possible to control not only when a particular type of DNA polymer forms or dissolves, but also the polymer composition, which depends on the level of active tiles at a particular point in time. To interconnect genes and control RNA levels over time we proposed a strategy relying on promoter “nicking” and strand displacement,^20,37^ however we expect that other approaches may be effective in generating nucleic acid regulators.^16,42^ Further, RNA levels could be tuned more finely by introducing RNA-degrading enzymes.^43^ Finally, our strategy takes advantage of simple DNA gates to route a single RNA output to control two processes simultaneously: the activation of DNA tiles, and the activation of downstream genes. We expect this approach could be expanded so that a single gene could control multiple processes in parallel, while maintaining modularity and minimizing the need to design additional genes.

Our efforts take advantage of the versatility and composability of nucleic acid components, which have made it possible to build rapidly growing libraries of circuits^44^ and structures^19^. New exciting demonstrations of autonomous biomaterials have emerged from the integration of circuits generating temporal signals and of self-assembling elements with the capacity to sense and process such signals.^7,8,45^ Here we have used genetic cascades to build an autonomous system of multiple DNA nanotubes, a remarkably simple DNA nanostructure, but we expect that they could be repurposed to coordinate the assembly of very different structural parts, like DNA origami,^46,47^ nanoparticles,^48,49^ and coacervates^50^. Overall, our work suggests a way toward scaling up the complexity of biomolecular materials by taking advantage of the timing of “molecular instructions” for self-assembly, rather than by increasing the number of molecules carrying such instructions. This points to the exciting possibility of generating distinct materials that can spontaneously “develop” from the same finite set of parts, by simply rewiring the elements that control the temporal order of assembly.

## Acknowledgements

Experimental work at UCLA was supported by the U.S. Department of Energy (DOE), Office of Science, Basic Energy Sciences (BES) under Award # DE-SC0010595 to EF. Computational modeling work was supported by the National Science Foundation (NSF) under CCF award number # 2107483 to EF. This work was supported by the European Research Council, ERC (project n.819160) (FR), by Associazione Italiana per la Ricerca sul Cancro, AIRC (project n. 21965) (FR), by the Italian Ministry of University and Research (Project of National Interest, PRIN,2017YER72K).

