## Supplementary Information for "Developmental assembly of multi-component polymer systems through interconnected gene networks *in vitro*"

| | |
|---|---|
| **Materials** | **2** |
| **Methods** | **9** |
| **Supplementary Figures** | **12** |
| **Supplementary Note 1: Modeling** | **26** |
| **References** | **29** |

### Materials

#### Chemicals

Reagent-grade chemicals (Trizma base (cat. #93352), acetic acid (cat. #A6283), boric acid (cat. #B0252), Ethylenediaminetetraacetic acid (EDTA) (cat. #798681), acrylamide-bis-acrylamide (40%) (cat. #A7168), ammonium persulfate (APS) (cat. #A7460), and N,N,N$^{I}$,N$^{I}$-tetramethyl ethylenediamine (TEMED) (cat. #T9281)) were purchased from Sigma-Aldrich (St Louis, Missouri) and used without further purification. The ORangeRuler 50 bp DNA ladder was purchased from ThermoFisher Scientific (USA) (cat. #SM0613). Transcription buffer was purchased from ThermoFisher Scientific (USA) and stored at -20°C. 5X transcription buffer contains: 200 mM Tris-HCl, 30 mM $MgCl_2$, 50 mM DTT, 50 mM NaCl, and 10 mM spermidine (pH 8.0 at 25°C). Nucleotide triphosphates (NTPs) were purchased from Jena Bioscience (Germany) and stored at -20°C.

#### Enzymes

T7 RNA polymerase was purchased from ThermoFisher Scientific (USA) ((cat. #EP0113). The bulk batch of T7 RNA polymerase was split into smaller aliquots (enough for 20–30 experiments) to minimize degradation of the enzyme by repeated removal from the freezer, and stored at -20°C.

#### Oligonucleotides

HPLC-purified oligonucleotides were purchased from IDT DNA Technologies (Coralville, IA) and Metabion International AG (Planegg, Germany). All oligonucleotides were dissolved in double-distilled $H_2O$ at a concentration of 100 μM and aliquoted at -20°C for long-term storage. All sequences were designed using Nupack or IDT software programs.[1,2] The concentration of oligonucleotides was quantified by UV-vis spectrophotometry using a Thermo Fisher Scientific Multiskan SkyHigh Microplate Spectrophotometer.

#### Tile designs and assembly

Building on previous work, we have re-engineered three different DNA tiles (green, red, and blue) that contain the same 5-nt sticky ends portion responsible for self-assembly but differ in the toehold sequence that can be orthogonally targeted by different regulatory strands.[3,4] A stock solution of DNA tiles was first formed from five strands (referred to as S1-S5) at a concentration of 5 μM and annealed with a Bio-Rad Mastercycler Gradient Thermocycler by heating to 90 °C and cooling to 25 °C at a constant rate over a period of 6 hours.

In addition to the DNA sequences, this section also includes schematic representations of each tile type. Nucleotides in *italics* for strands S2 and S4 denote the sticky end portions. Strand S3 was conjugated to a fluorophore at the 5' end. Green, red, or blue circles on the blue strand (S3) represent the position of the Cy3, Cy5, or 6- FAM fluorophores, as indicated in the strand sequence. Strand S2 also contains the 7-nt inhibitor binding domain (**bold**). The inhibitor strand binds to S2 via a 14-nt portion that first binds to the 7-nt inhibitor-binding domain of S2 and then invades the 5-nt sticky end and 2 additional nucleotides. The inhibitor strand contains an additional 6-nt portion (<u>underlined</u>) that remains available for binding of the activator strand. The activator first binds to this portion and then invades the duplex formed from the inhibitor and S2 strand, displacing the inhibitor from the tile and restoring the tile's ability to self-assemble.

##### Green system: Tile 1

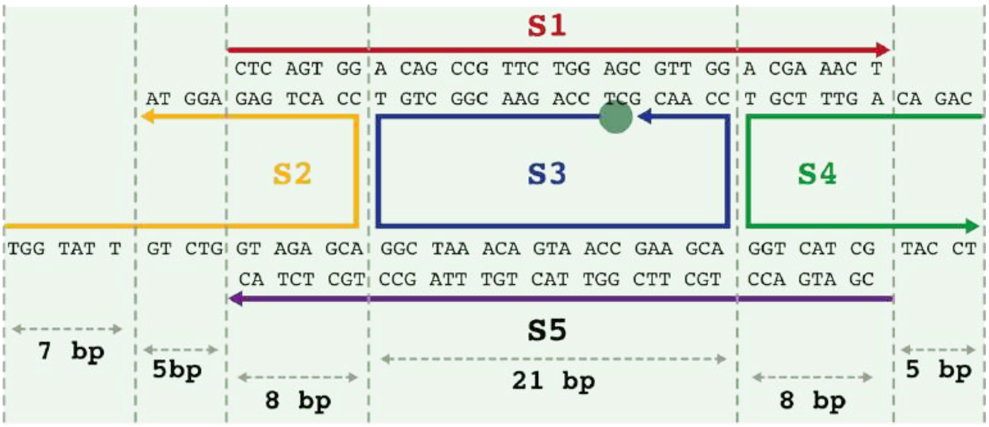


***Scheme S1.*** *5-nucleotide sticky end green tile system. DNA tiles consist of 5 ssDNA (S1-S5). Here each strand is highlighted with a different color. The arrow indicates the 3' end.*

| **Name** | **Sequence** |
|---|---|
| S1 | 5'- CTC AGT GGA CAG CCG TTC TGG AGC GTT GGA CGA AAC T |
| S2 | 5'- **TGG TAT T** *GTC TG* GTA GAG CAC CAC TGA G *AGG TA* |
| S3 (G) | 5' - (Cy3) -TCC AGA ACG GCT GTG GCT AAA CAG TAA CCG AAG CAC CAA CGC T |
| S4 | 5' - *CAGAC* AG TTT CGT GGT CAT CGT ACC T |
| S5 | 5' - CGA TGA CCT GCT TCG GTT ACT GTT TAG CCT GCT CTA C |
| DNA Inhibitor (G) | 5' - ACC AGA CAA TAC CA ATC CGC |
| RNA Inhibitor (G) | 5' - ACC AGA CAA UAC CA AUC CGC |
| DNA Activator (G) | 5'- GCG GAT TGG TAT TGT CTG GT |
| RNA Activator (G) | 5'- GCG GAU UGG UAU UGU CUG GU |

#### Red system: Tile 2

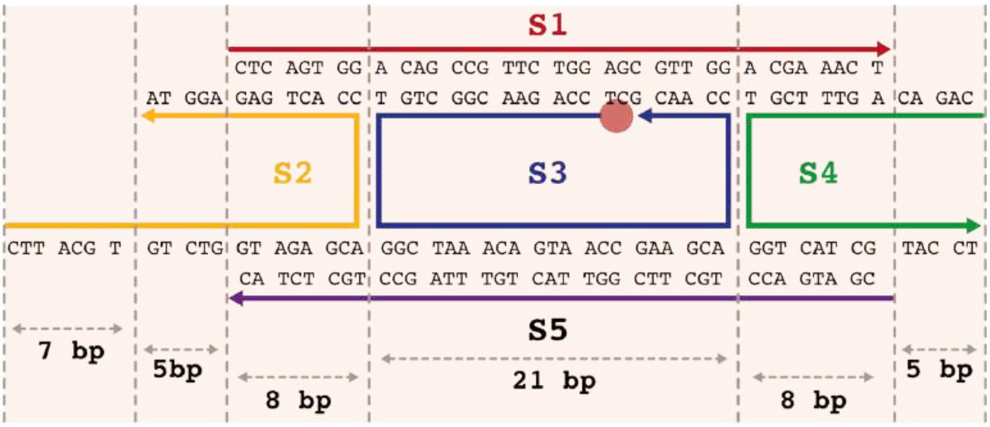


***Scheme S2.*** *5-nucleotide sticky end red tile system. DNA tiles consist of 5 ssDNA (S1-S5). Here each strand is highlighted with a different color. The arrow indicates the 3' end.*

| **Name** | **Sequence** |
|---|---|
| S1 | 5'- CTC AGT GGA CAG CCG TTC TGG AGC GTT GGA CGA AAC T |
| S2 | 5'- **CTT ACG T** *GTC TG* GTA GAG CAC CAC TGA G *AGG TA* |
| S3 (R) | 5' - (Cy5) -TCC AGA ACG GCT GTG GCT AAA CAG TAA CCG AAG CAC CAA CGC T |
| S4 | 5' - *CAGAC* AG TTT CGT GGT CAT CGT ACC T |
| S5 | 5' - CGA TGA CCT GCT TCG GTT ACT GTT TAG CCT GCT CTA C |
| DNA Inhibitor (R) | 5' - ACC AGA CAC GTA AGG ATG GC |
| RNA Inhibitor (R) | 5' - ACC AGA CAC GUA AGG AUG GC |
| DNA Activator (R) | 5'- GCC ATC CTT ACG TGT CTG GT |
| RNA Activator (R) | 5'- GCC AUC CUU ACG UGU CUG GU |

**Blue system: Tile 3**

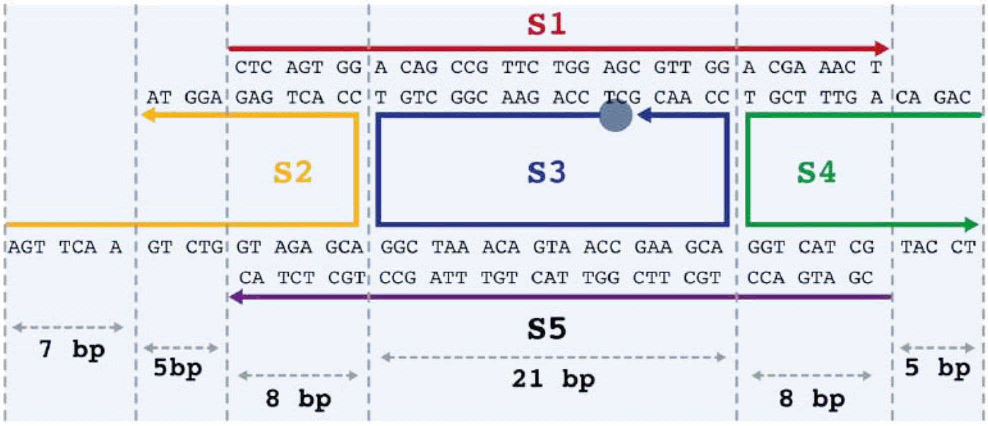


***Scheme S3.*** *5-nucleotide sticky end blue tile system. DNA tiles consist of 5 ssDNA (S1-S5). Here each strand is highlighted with a different color. The arrow indicates the 3' end.*

| **Name** | **Sequence** |
|---|---|
| S1 | 5'- CTC AGT GGA CAG CCG TTC TGG AGC GTT GGA CGA AAC T |
| S2 | 5'- **AGT TCA A** *GTC TG* GTA GAG CAC CAC TGA G *AGG TA* |
| S3 (B) | 5' - (6-FAM) -TCC AGA ACG GCT GTG GCT AAA CAG TAA CCG AAG CAC CAA CGC T |
| S4 | 5' - *CAGAC* AG TTT CGT GGT CAT CGT ACC T |
| S5 | 5' - CGA TGA CCT GCT TCG GTT ACT GTT TAG CCT GCT CTA C |
| DNA Inhibitor (B) | 5' - ACC AGA CTT GAA CTT CGA CC |
| RNA Inhibitor (B) | 5' - ACC AGA CUU GAA CUU CGA CC |
| DNA Activator (B) | 5'- GGT CGA AGT TCA AGT CTG GT |
| RNA Activator (B) | 5'- GGU CGA AGU UCA AGU CUG GU |

#### Genes design

In the following sequences, the underlined portion represents the promoter site recognized by T7 RNA polymerase, while the *italics* domain represents the portion that is transcribed. The genes contain "sealing" domains in the genes at the 5' end of the non-template strand to prevent breathing at the promoter site. To ensure good transcriptional yield, each strand was designed to begin with G. Hence one **G** (**bold**) was added after the promoter sequence.

**Table 1: High yield design**

| **Gene** | **Sequence** |
|---|---|
| NonTemplate Activator 1 | 5'- G CGC TAA TAC GAC TCA CTA TA **G** *GCG GAT TGG TAT TGT CTG GT* |

| | |
|---|---|
| Template Activator 1 | 5'- ACC AGA CAA TAC CAA TCC GCC TAT AGT GAG TCG TAT TAG CGC |
| NonTemplate Activator 2 | 5'- CAA TAC CAA TCC GC TAA TAC GAC TCA CTA TA **G** *GCC ATC CTT ACG TGT CTG GT* |
| Template Activator 2 | 5' - ACC AGA CAC GTA AGG ATG GCC TAT AGT GAG TCG |
| NonTemplate Activator 3 | 5' – CAC GTA AGG ATG GC TAA TAC GAC TCA CTA TA **G** *GGT CGA AGT TCA AGT CTG GT* |
| Template Activator 3 | 5' - ACC AGA CTT GAA CTT CGA CCC TAT AGT GAG TCG |
| NonTemplate Inhibitor 1 | 5'- CT TGA ACT TCG ACC TAA TAC GAC TCA CTA TA **G** *ACC AGA CAA TAC CA ATC CGC* |
| Template Inhibitor 1 | 5'- GCG GAT TGG TAT TGT CTG GTC TAT AGT GAG TCG |
| NonTemplate Inhibitor 2 | 5'- TGG TAT TGT CTG GT TAA TAC GAC TCA CTA TA **G** *ACC AGA CAC GTA AGG ATG GC* |
| Template Inhibitor 2 | 5'- GCC ATC CTT ACG TGT CTG GTC TAT AGT GAG TCG |
| NonTemplate Inhibitor 3 | 5'- CTT ACG TGT CTG GT TAA TAC GAC TCA CTA TA **G** *ACC AGA CTT CGA CC* |
| Template Inhibitor 3 | 5'- GGT CGA AGT CTG GTC TAT AGT GAG TCG |

**Table 2. Connector system for high yield design**

| **Connector system** | **Sequence** |
|---|---|
| Released Strand 1_6nt Toehold | 5'- TAA TA GCG TGG TAT TG |
| Non Released Strand 1 | 5'- ACC AGA CA ATA CCA ATC CGC |
| Released Strand 2_6nt Toehold | 5'- TAT TA GCC ATC CTT ACG TG |
| Non Released Strand 2 | 5' – ACC AGA CA CGT AAG GAT GGC |
| Released Strand 3_6nt Toehold | 5'- TAT TA GGT CGA AGT TCA AG |
| Released Strand 3_5nt Toehold | 5'- TAT TA GGT CGA AGT TCA AGT |
| Released Strand 3_4nt Toehold | 5'- TAT TA GGT CGA AGT TCA AGT C |
| Released Strand 3_3nt Toehold | 5'- TAT TA GGT CGA AGT TCA AGT CT |

| | |
|---|---|
| Non Released Strand 3 | 5' - ACC AGA CT TGA ACT TCG ACC |
| Released Strand 4_6nt Toehold | 5'- TAT TA ACC AGA CAA TAC CA |
| Released Strand 4_ 5nt Toehold | 5'- TAT TA ACC AGA CAA TAC CAA |
| Released Strand 4_4nt Toehold | 5'- TAT TA ACC AGA CAA TAC CAA T |
| Released Strand 4_3nt Toehold | 5'- TAT TA ACC AGA CAA TAC CAA TC |
| Non Released Strand 4 | 5'- GCG GAT TG GTA GTT TCT GGT |
| Released Strand 5_6nt Toehold | 5'- TAT TA ACC AGA CAC GTA AG |
| Release Strand 5_5nt Toehold | 5'- TAT TA ACC AGA CAC GTA AGG |
| Released Strand 5_4nt Toehold | 5'- TAT TA ACC AGA CAC GTA AGG A |
| Released Strand 5_3nt Toehold | 5'- TAT TA ACC AGA CAC GTA AGG AT |
| Non Released Strand 5 | 5'- GCC ATC CT TAC GTG TCT GGT |

**Table 3. Modified sequences for high yield design**

To study the strand displacement reaction that occurs after the RNA activator is produced during gene transcription, we used the following sequences modified with fluorophores and quenchers.

| **Modified strands** | **Sequence** |
|---|---|
| Released Strand 1 | 5'- TAT TA GCG TGG TAT TG - (Cy3) |
| Non Released Strand 1 | 5'- (BHQ-1) - ACC AGA CA ATA CCA ATC CGC |
| Released Strand 2 | 5'- TAT TA GCC ATC CTT ACG TG - (Cy5) |
| Non Released Strand 2 | 5'- (BHQ-2)- ACC AGA CA CGT AAG GAT GGC |
| S2 (G)_Q | 5'- TGG TAT T GTC TG GT – (BHQ-1) - A GAG CAC CAC TGA G AGG TA |
| S2 (R)_Q | 5'- CTT ACG TGTC TG GT – (BHQ-2) – A GAG CAC CAC TGA G A GG TA |

| | |
|---|---|
| RNA Inhibitor (G)_F | 5'- (Cy3) - ACC AGA CAA UAC CA AUC CGC |
| RNA Inhibitor (R)_F | 5'- (Cy5) - ACC AGA CAC GUA AGG AUG GC |

**Table 4: Low yield design**

| **Gene** | **Sequence** |
|---|---|
| NonTemplate Activator 1 | 5'- G CGC <u>TAA TAC GAC TCA CTA TA</u> **G** *GCG GAT TGG TAT TGT CTG GT* |
| Template Activator 1 | 5'- ACC AGA CAA TAC CAA TCC GCC TAT AGT GAG TCG TAT TAG CGC |
| NonTemplate Activator 2 | 5'- <u>C GAC TCA CTA TA</u> **G** *GCC ATC CTT ACG TGT CTG GT* |
| Template Activator 2 | 5'- ACC AGA CAC GTA AGG ATG GCC TAT AGT GAG TCG GTA TTA CCA GAC AAT ACC A |
| NonTemplate Activator 3 | 5'- <u>C GAC TCA CTA TA</u> **G** *GGT CGA AGT TCA AGT CTG GT* |
| NonTemplate 3 | 5'- AC CAG ACT TGA ACT TCG ACC CTA TAG TGA GTC GTA TTA A CCA GAC ACG TAA G |

**Table 5. Connector system for low yield design**

| **Connector system** | **Sequence** |
|---|---|
| Released Strand 1 | 5'- TGG TAT TGT CTG GT TAA TA |
| Non Released Strand 1 | 5'- ACC AGA CA ATA CCA ATC CGC |
| Released Strand 2 | 5'- CTT ACG TGT CTG GT TAA TA |
| Non Released Strand 2 | 5' – ACC AGA CA CGT AAG GAT GGC |

**Table 6. Modified sequences for low yield design**

| **Modified strands** | **Sequence** |
|---|---|
| Released Strand 1 | 5'- TGG TAT TGT CTG GT TAA TA - (Cy3) |
| Non Released Strand 1 | 5'- (BHQ-1) - ACC AGA CA ATA CCA ATC CGC |
| Released Strand 2 | 5'- CTT ACG TGT CTG GT TAA TA - (Cy5) |
| Non Released Strand 2 | 5'- (BHQ-2)- ACC AGA CA CGT AAG GAT GGC |

### Methods

#### Fluorescence measurements

All fluorescence experiments were performed at 30° C, in 1X transcription buffer, 10 mM NTPs, pH 8.0 in a 100 µL cuvette (total volume of solution 100 µL). Equilibrium fluorescence measurements were performed using a Cary Eclipse fluorimeter. Excitation was at 550 (±5) nm and acquisition at 570 (±5) nm (for strands labeled with Cy3) or at 650 (±5) nm and acquisition at 670 (±5) nm (for strands labeled with Cy5).

#### Fluorescence measurements for tile activation

The self-assembly of DNA tiles driven by RNA activator transcription was studied by first examining the assembly of individual tiles (1, and 2) at a fixed concentration of RNA inhibitor strand (1, and 2, 1 µM) in the presence of different concentrations of the transcribing gene for the RNA activator strand and the T7 RNA polymerase for both strategies described in the main text. Similarly, to study the assembly of DNA nanotubes driven by green and red activators, equimolar concentrations of inactive tiles 1 and 2 (250 nM labeled tiles/1 µM RNA inhibitor) were mixed in the absence or presence of the connector system (300 nM) in the same buffer solution at different concentrations of the transcribing genes for the RNA activator strands and the T7 RNA polymerase enzyme. The percentage of tile activation, and thus inhibitor release, was tracked with inactive tiles labeled with a fluorophore/quencher pair so that tile activation could be tracked by the increase in fluorescence signal due to inhibitor release. We recorded the fluorescence signal in real time until it reached equilibrium before the addition of the T7 RNA polymerase enzyme.

#### Data analysis for tile activation

The values for RNA inhibitor released from tiles (nM) are calculated from the relative fluorescence signal registered upon addition of a saturating concentration of T7 RNA polymerase (4 U/µL) in the presence of the specific activator gene according to the following formula:

$$[Released\ Inhibitor] = \frac{[Inhibitor] * (F_{+T7\ RNAP} - F_{-T7\ RNAP})}{F_{max}}$$

where $F_{+T7\ RNAP}$ is the fluorescence signal observed after the addition of T7 RNA polymerase in the presence of a fixed concentration of activator gene; $F_{-T7\ RNAP}$ is the fluorescence signal observed in the absence of T7 RNA polymerase at a fixed concentration of activator gene; $F_{max}$ is the maximum fluorescence value that can be achieved when 250 nM of inhibitor is released from the DNA tiles; [Inhibitor] is 250 nM and is the maximum concentration that can be released from the DNA tiles (250 nM).

#### Active tile preparation

DNA tiles for all systems presented were prepared as follows. The solution of active tiles was prepared to a target concentration of 5 µM by mixing S1-S5 strands of each tile type (1, 2, and 3) in stoichiometric ratios with 1X transcription buffer and double distilled water ($ddH_2O$) in DNA Lo-bind tubes. The solution was annealed using an Eppendorf Mastercycler Gradient Thermocycler by heating to 90 °C and cooling to 25 °C at a constant rate for a period of 6 hours. The annealed tiles were then diluted to a final concentration of 250 nM in

1X transcription buffer. The solution of the assembled tiles was diluted to 50 nM in the same buffer before a microscope image was taken.

##### Inactive tile preparation

The annealed tiles (1, 2, and 3) were diluted in 1X transcription buffer to a final concentration of 250 nM each, and incubated in the Mastercycler at 30 °C for 1 h prior to adding the RNA inhibitor strand. The assembled tiles solution was diluted to 50 nM in the same buffer before a microscope image was taken. The inhibitor strands for tiles 1, 2, and 3 (G, R and B) were then added to an excess concentration of 1 μM to ensure the complete disassembly of all DNA tiles. The reaction was allowed to proceed for 30 minutes at 30 °C and then we imaged the samples using a fluorescence microscope.

##### Reassembly by addition of synthetic RNA activator strand

The annealed tiles were diluted to a final concentration of each tile of 250 nM. The RNA inhibitor 1 (G) was then added at a concentration of 1 μM and the microscope image was taken after 30 minutes. The RNA inhibitor 2 (R) was added to a separate solution at a concentration of 1 μM and the microscopy image was taken after 30 minutes. The RNA inhibitor 3 (B) was added to a separate solution at a concentration of 1 μM and the microscopy image was taken after 30 minutes. The RNA activators 1, 2 and 3 (G, R, B) were then respectively added at a concentration of 3 μM and the microscope image was taken for the next 24 hours. The same protocol was followed for mixed G/R/B nanotubes reassembly. The RNA inhibitors 1, 2 and 3 (G, R and B) were added at a concentration of 1 μM to the annealed tiles (G, R and B), respectively. Samples were incubated at 30°C and a confocal image was taken after 1 hour. Of note, the tiles disassemble. The RNA activators 1, 2, and 3 (G, R, B) were then added at a concentration of 3 μM and the confocal images were taken in the next 24 hours.

##### Reassembly via transcription of RNA activator strand

Inactive tiles 1, 2, and 3 (G, R, and B, respectively) were incubated at 30°C for 1 hour before the addition of T7 RNA polymerase to produce the RNA activator strand. The annealed synthetic gene for transcribing the corresponding RNA activator strand (1, 2, and 3) was mixed with the inactive tiles (250 nM each) and transcription mix (T7 RNAP 4U/μL, 1X transcription buffer, 10 mM each nucleoside triphosphates (NTPs)) at 30 °C and observed under a fluorescence microscope for several hours.

##### Disassembly and assembly via transcription of RNA regulators

Inactive tiles G, R, and B were incubated at 30 °C for 1 h prior to production of the RNA activator strand. The annealed synthetic gene for transcribing the corresponding RNA activator (G, R, B) and inhibitor strands were mixed with the inactive tiles (250 nM each), connector complexes (300 nM) and transcription mix (T7 RNAP, 1X transcription buffer, 10 mM each nucleoside triphosphates (NTPs)) at 30 °C and observed under a fluorescence microscope for several hours.

##### Fluorescence Microscopy experiments

Fluorescence imaging of DNA nanostructures for fluorescence microscopy, the central strand of each tile (S3) was labeled at the 5’ end with a different fluorophore (Cy3, Cy5, or 6-FAM). Nanotube samples were imaged using an inverted microscope (Zeiss Observer 7) with a 100× oil immersion objective (EC Plan-Neo Fluor) and a monochrome Axiocam 305 camera. For imaging, samples were diluted with the experimental buffer to a final concentration of 50 nM of each tile. A 5 µL drop of this diluted solution was then deposited between clean microscope coverslips (Menzel-Glaser; thickness: 0.13 – 0.16 mm; size: 18 x 18 mm). Images were acquired using 90 HE LED filters (EX: 385, 475, 555, 630 nm; QBS 405 + 493 + 575 + 653; EM: QBP 425/30 + 514/30 +592/30 + 709/100). The exposure time was set to 10000 ms. Ten images at the same location were processed to correct for uneven illumination and superimposed to produce multicolor images using ZEN 3.6 software (ZEISS).

##### Fluorescence microscopy data and image processing

Fluorescence microscopy images were analyzed and processed using ZEN 3.6 software (ZEISS) to correct for uneven illumination and overlaid to produce multicolor images. Branched or looping nanotubes were removed from the length dataset using an in-house MATLAB script. ImageJ was used to analyze the pixel intensity of the selected structure for each individual channel. Colocalization analysis was performed using a plugin for ImageJ software called JACoP (Just Another Co-localization Plugin).[5,6]

##### Native PAGE experiments

Native polyacrylamide gels (15%) were prepared with TBE (10X), ammonium persulfate (APS), and tetramethylethylenediamine (TEMED) at appropriate ratios. Gels were cast in 10 × 10 cm, 1.5 mm thick disposable minigel cassettes and allowed to polymerize for at least 30 min before electrophoresis. The gel was incubated with a running buffer (1X TBE solution, pH 8.0) for 30 minutes at 25°C. Sample volumes of 10 µl were combined with 1 µl of 6× Orange DNA Loading Dye and then loaded directly into the gel. Native PAGE was run in a mini-PROTEAN tetracell electrophoresis unit (Bio-Rad) at 25°C using 1X TBE buffer at pH 8.0 and a constant voltage of 110 V for 2 h 30 min (using the Bio-Rad PowerPac Basic power supply). Gels were stained with SYBR Gold and scanned using a ChemiDoc MP imaging system (Bio-Rad).

##### Statistical Analysis

All data shown in graphs are presented as mean ± standard deviation (SD). Statistical analysis was performed using GraphPad Prism 8 software. No specific preprocessing of data was performed prior to statistical analyses.

#### Supplementary Figures

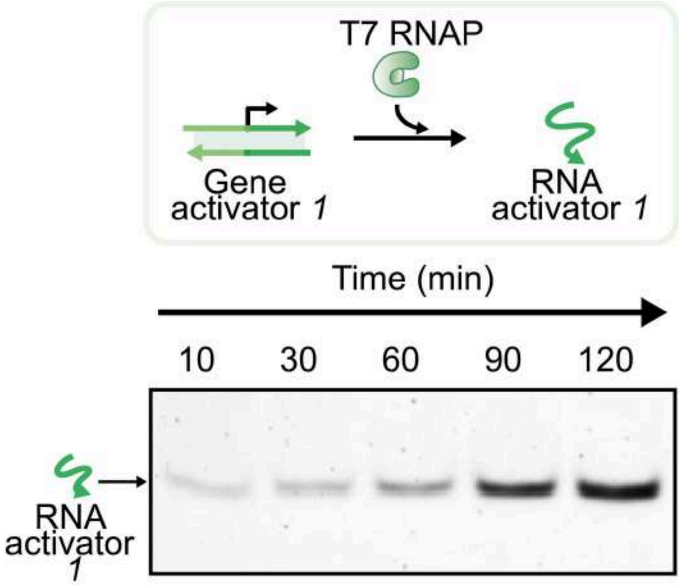


***Supplementary Figure 1.*** *Native PAGE gel (15% polyacrylamide) showing production of RNA activator 1 over time. Each solution containing gene activator 1 (100 nM) was prepared in a 1X transcription buffer, NTPs (10 mM), T7 RNAP (4 U/μL). Gel was run at 25°C (110 V) for 2h 30 min in 1x TBE buffer, pH 8.0, stained with 1x SYBR-Gold, and imaged using the Gel Doc XR system (Bio-Rad).*

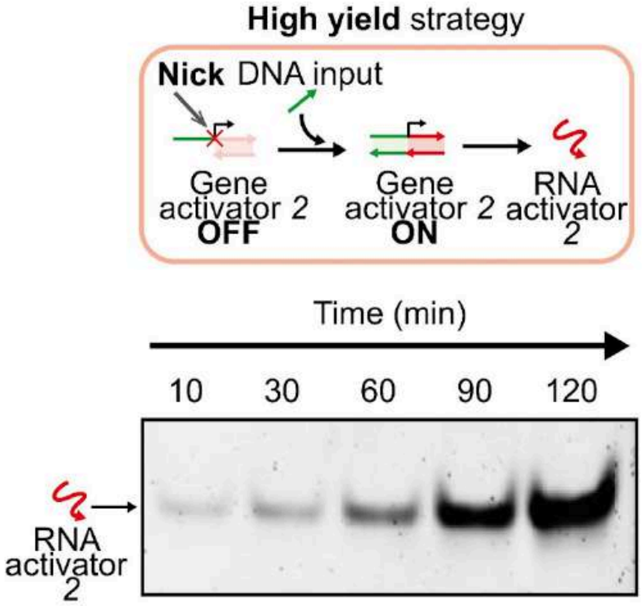


***Supplementary Figure 2.*** *Native PAGE gel (15% polyacrylamide) showing production of RNA activator 2 from a high yield gene over time. Each solution containing high yield gene 2 (100 nM), and DNA input (100 nM) was prepared in a 1X transcription buffer, NTPs (10 mM), T7 RNAP (4 U/μL). Gel was run at 25°C (110 V) for 2h 30 min in 1x TBE buffer, pH 8.0, stained with 1x SYBR-Gold, and imaged using the Gel Doc XR system (Bio-Rad).*

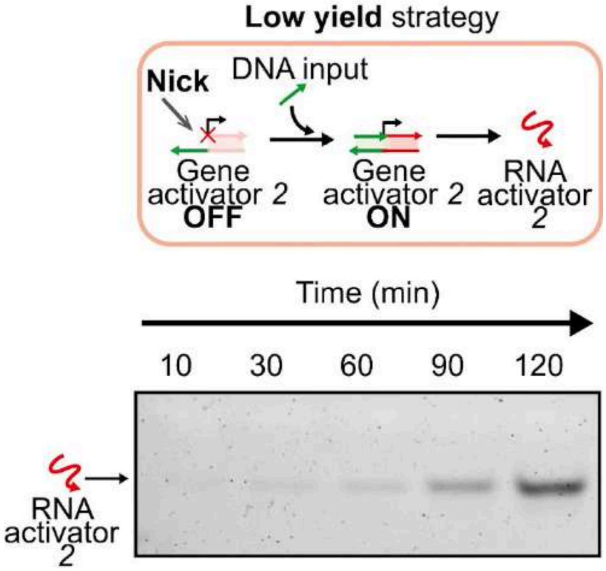


***Supplementary Figure 3.*** *Native PAGE gel (15% polyacrylamide) showing production of RNA activator 2 from a low yield gene over time. Each solution containing low yield gene 2 (100 nM), and DNA input (100 nM) was prepared in a 1X transcription buffer, NTPs (10 mM), T7 RNAP (4 U/µL). Gel was run at 25°C (110 V) for 2h 30 min in 1x TBE buffer, pH 8.0, stained with 1x SYBR-Gold, and imaged using the Gel Doc XR system (Bio-Rad).*

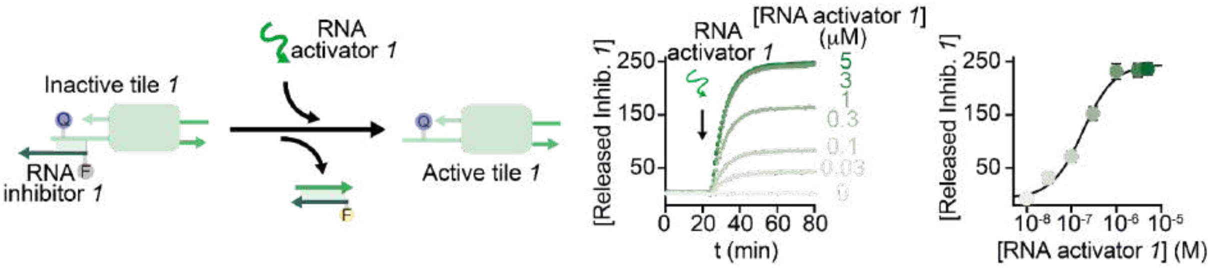


***Supplementary Figure 4. Left.*** *Tile 1 activation using synthetic RNA activator 1.* ***Middle.*** *Released inhibitor 1 at different concentrations of synthetic RNA activator 1 in the presence of tile 1 (250 nM) and RNA inhibitor 1 (1 µM).* ***Right.*** *End point values vs RNA activator 1 concentration. Experiments were performed at 30° C in 1X transcription buffer, 10 mM NTPs, pH 8.0 in a 100 µL cuvette. Experimental values are averages of three separate measurements and error bars reflect standard deviations.*

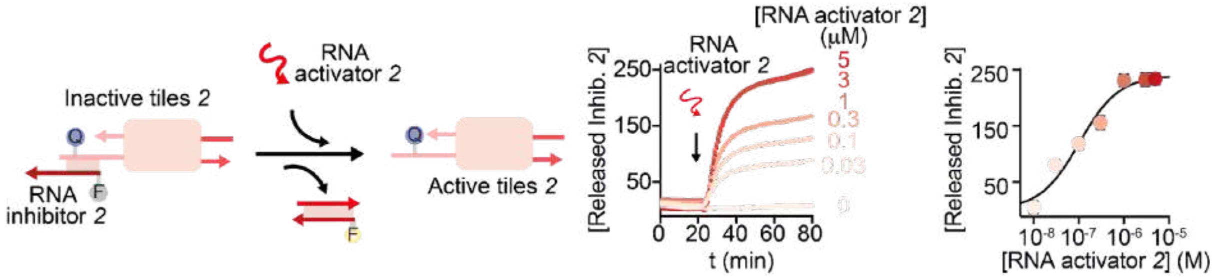


***Supplementary Figure 5. Left.*** *Tile 2 activation using synthetic RNA activator 2.* ***Middle.*** *Released inhibitor 2 at different concentrations of synthetic RNA activator 2 in the presence of tile 2 (250 nM) and RNA inhibitor 2 (1 µM).* ***Right.*** *End point values vs RNA activator 2 concentration. Experiments were performed at 30° C, in 1X transcription buffer, 10 mM NTPs, pH 8.0 in a 100 µL cuvette. Experimental values are averages of three separate measurements and error bars reflect standard deviations.*

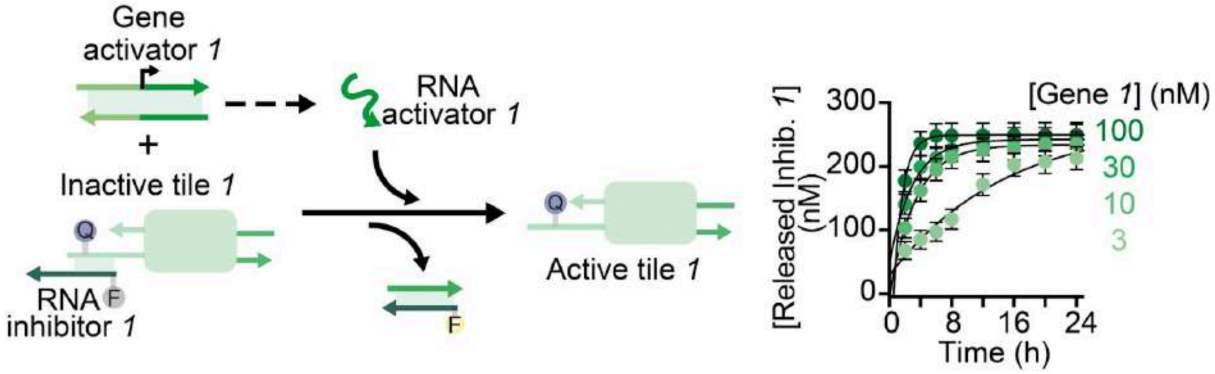


***Supplementary Figure 6. Left.*** *Tile 1 (250 nM) activation using gene activator 1 (100 nM).* ***Right.*** *Kinetic traces showing the release of the RNA inhibitor over time at different concentrations of gene activator 1. Experiments were performed at 30° C in 1X transcription buffer, 10 mM NTPs, T7 RNP 4 U/µL, pH 8.0 in a 100 µL cuvette. Experimental values are averages of three separate measurements and error bars reflect standard deviations.*

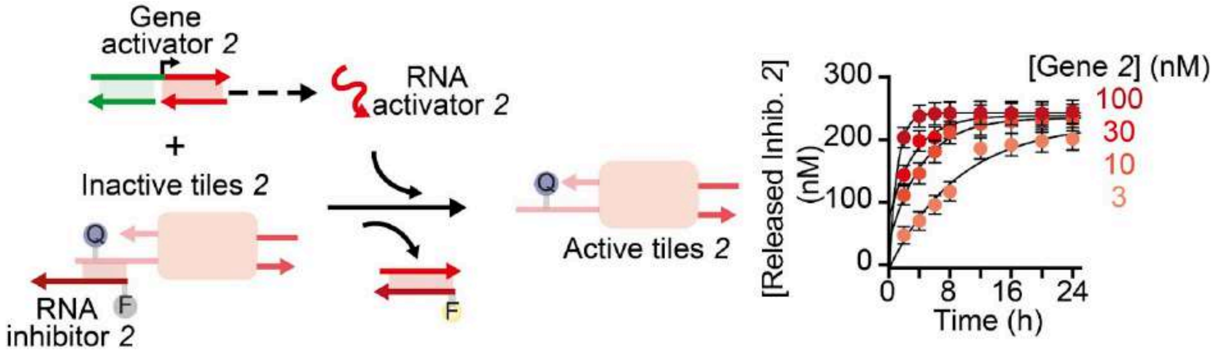


***Supplementary Figure 7. Left.*** *Tile 2 (250 nM) activation using gene activator 2 (100 nM).* ***Right.*** *Kinetic traces showing the release of the RNA inhibitor 2 over time at different concentrations of high yield gene activator 2. Experiments were performed at 30° C in 1X transcription buffer, 10 mM NTPs, T7 RNP 4 U/ uL, pH 8.0 in a 100 µL cuvette. Experimental values are averages of three separate measurements and error bars reflect standard deviations.*

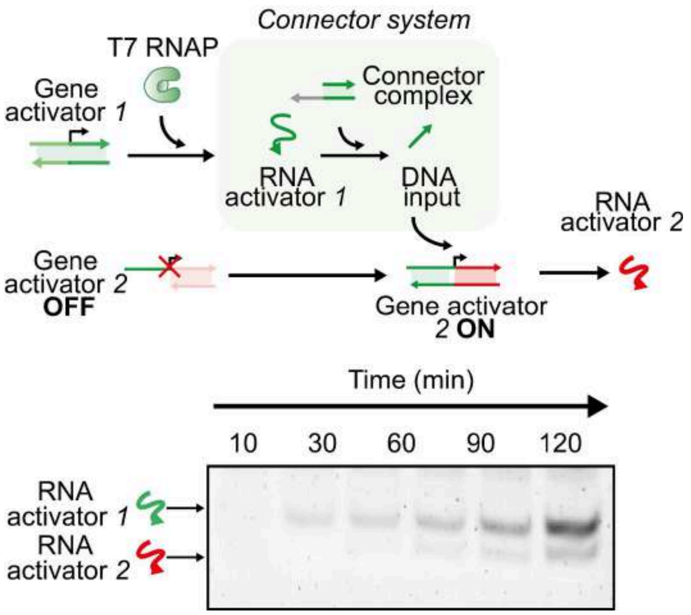


***Supplementary Figure 8. Top.*** *Schematic representation of a simplified cascade network of two interconnected synthetic genes (1 and 2).* ***Bottom.*** *Native PAGE gel (15% polyacrylamide) showing production of RNA activator 1 and 2 over time. We achieve a tunable delay in the production of RNA activator 2 by confirming the gene is not switched to ON until its promoter is completed. Each solution containing gene activator 1 and 2 (100 nM), and connector complex (300 nM) was prepared in a 1X transcription buffer, NTPs (10 mM), T7 RNAP (4 U/µL). Gel was run at 25°C (110 V) for 2h 30 min in 1x TBE buffer, pH 8.0, stained with 1x SYBR-Gold, and imaged using the Gel Doc XR system (Bio-Rad).*

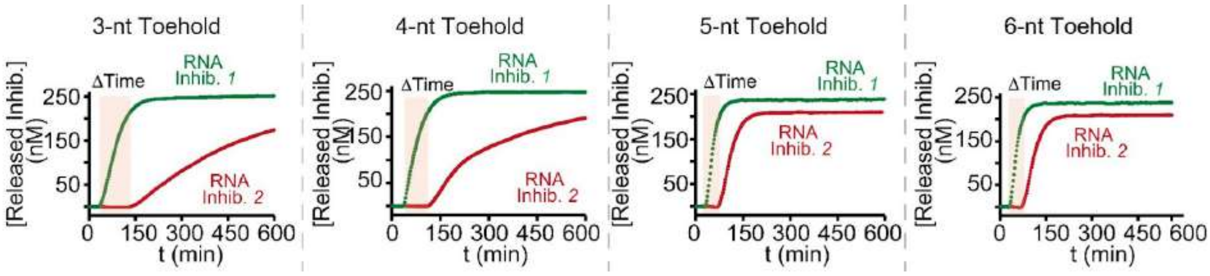


***Supplementary Figure 9.*** *Fluorescence kinetic traces showing the release of RNA inhibitor strands 1 and 2 from their corresponding tiles triggered by the presence of transcribed RNA activator strands 1 and 2 and the connector complex at different toehold length (from 3 to 6-nt). Each solution containing tile 1 and 2 (250 nM), RNA inhibitors 1 and 2 (1 µM each), gene 1 and high yield gene 2 (100 nM each), and the connector complex (300 nM) was prepared in a 1X transcription buffer, NTPs (10 mM), T7 RNAP (4 U/µL).*

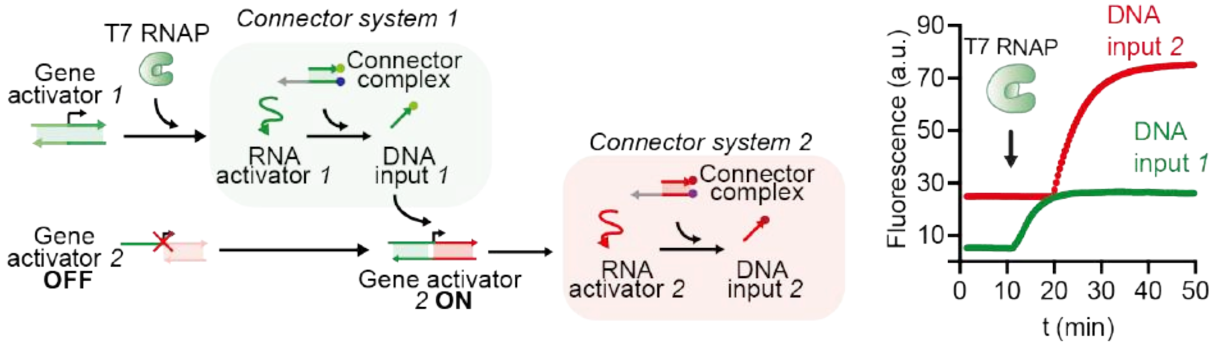


***Supplementary Figure 10. Left.*** *Schematic representation of the displacement of a DNA input from a connector complex upon RNA transcription.* ***Right.*** *Kinetic traces showing release of DNA input from fluorophore/quencher labeled connector complexes at fixed concentrations of gene 1 and high yield gene 2 (100 nM each). Each solution containing tile 1 and 2 (250 nM), RNA inhibitors (1 µM), gene activators 1 and 2 (100 nM), and the connector complexes (300 nM) was prepared in a 1X transcription buffer, NTPs (10 mM), T7 RNAP (4 U/µL).*

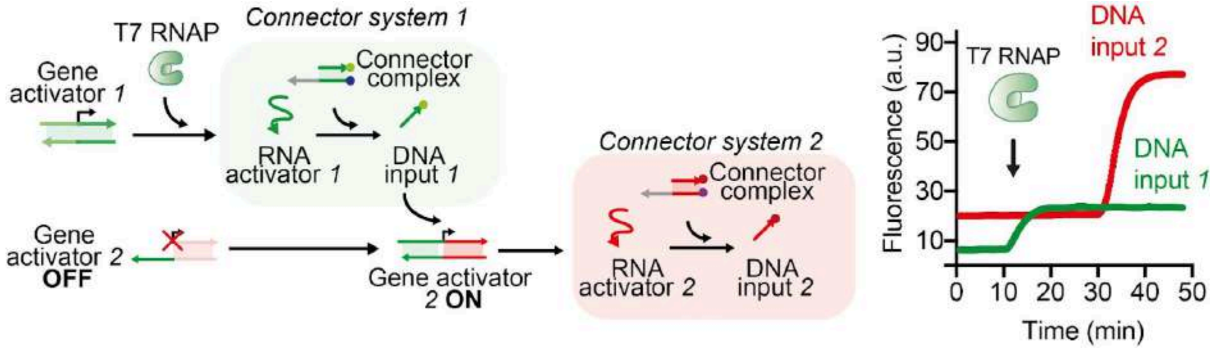


***Supplementary Figure 11.*** *Schematic representation of the displacement of a DNA input from a connector complex upon RNA transcription.* ***Right.*** *Kinetic traces showing release of DNA input from fluorophore/quencher labeled connector complexes at fixed concentrations of gene 1 and low yield yield gene 2 (100 nM each). Each solution containing tile 1 and 2 (250 nM), RNA inhibitors (1 µM), gene activators 1 and 2 (100 nM), and the connector complexes (300 nM) was prepared in a 1X transcription buffer, NTPs (10 mM), T7 RNAP (4 U/µL).*

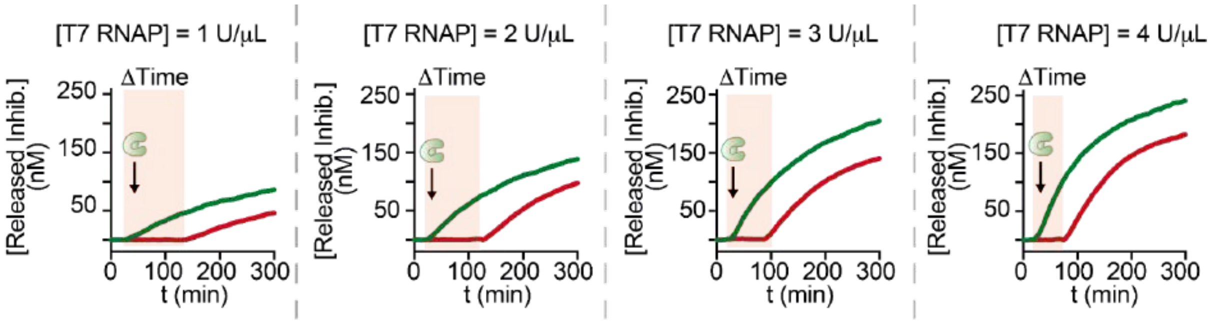


***Supplementary Figure 12.*** *Kinetic traces showing the release of RNA inhibitors (1 and 2) from the corresponding tiles at fixed concentrations of gene 1 and high yield gene 2 (100 nM each), connector complex (300 nM), and various concentrations of T7 RNAP (from 1 to 4 U/μL). To easily follow the release of the inhibitors after production of the RNA activator, we modified the RNA inhibitor and interacting tile strand with an appropriate fluorophore and quencher pair (1, Cy3/BHQ1 and 2, Cy5/BHQ2).*

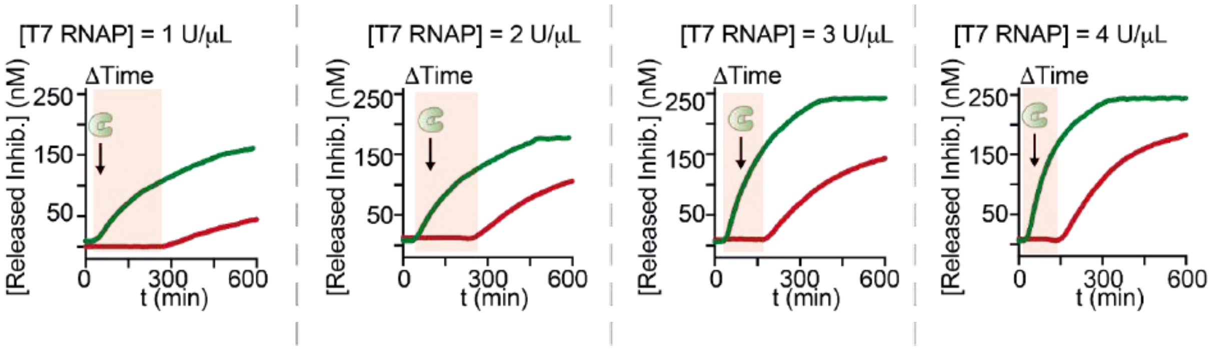


***Supplementary Figure 13.*** *Kinetic traces showing the release of RNA inhibitors (1 and 2) from the corresponding tiles at fixed concentrations of gene 1 and low yield gene 2 (100 nM each), connector complex (300 nM), and various concentrations of T7 RNAP (from 1 to 4 U/μL). To easily follow the release of the inhibitors after production of the RNA activator, we modified the RNA inhibitor and interacting tile strand with an appropriate fluorophore and quencher pair (1, Cy3/BHQ1 and 2, Cy5/BHQ2).*

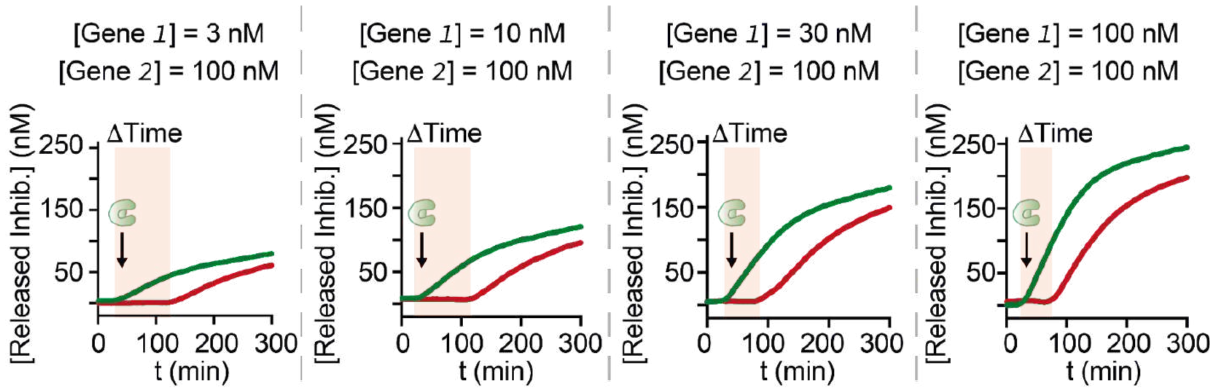


***Supplementary Figure 14.*** *Kinetic traces showing the release of RNA inhibitors (1 and 2) from the corresponding tiles at different concentrations of gene 1 (from 3 to 100 nM), fixed concentration of high yield gene 2 (100 nM), connector complex (300 nM), and T7 RNAP (4 U/uL). To easily follow the release of the inhibitors after the production of the RNA activator, we modified the RNA inhibitor and the interacting tile strand with a fluorophore and quencher pair (1, Cy3/BHQ1 and 2, Cy5/BHQ2).*

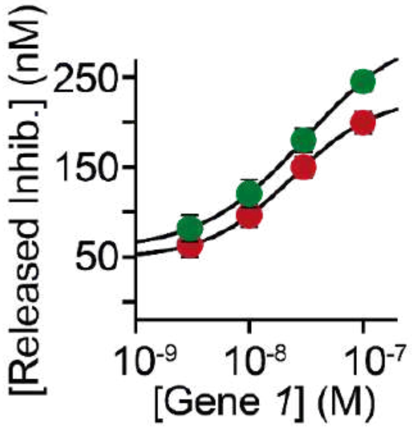


***Supplementary Figure 15.*** *Release of RNA inhibitors 1 (green) and 2 (red) from the corresponding DNA tiles after 300 min as a function of gene 1 concentration.*

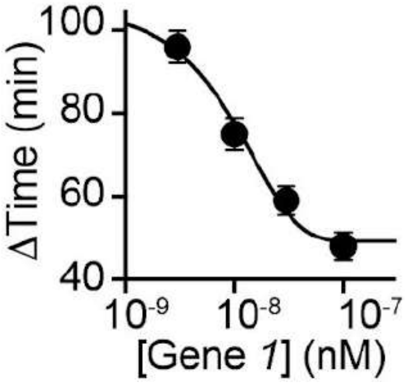


***Supplementary Figure 16.*** *Delay time (ΔT) of tile 2 activation as a function of gene 1 concentration.*

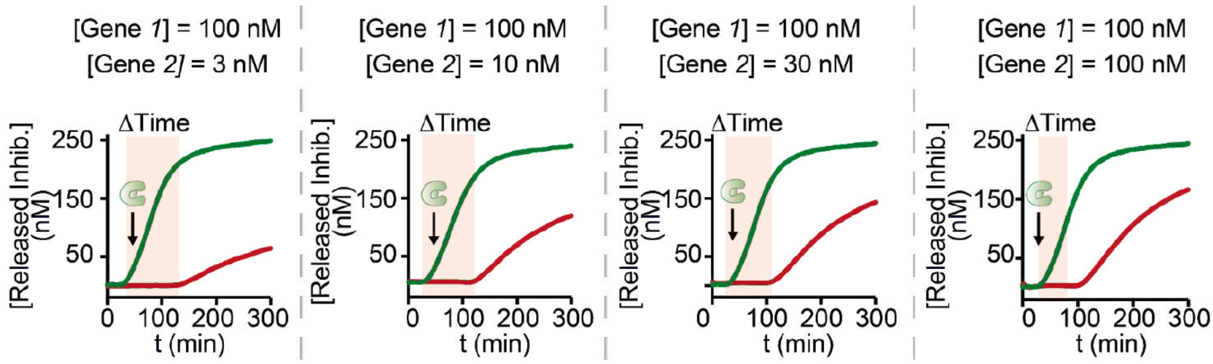


***Supplementary Figure 17.*** *Kinetic traces showing the release of RNA inhibitors (1 and 2) from the corresponding tiles at different concentrations of high yield gene 2 (from 3 to 100 nM), fixed concentration of gene 1 (100 nM), connector complex (300 nM), and T7 RNAP (4 U/μL). To easily follow the release of the inhibitors after the production of the RNA activator, we modified the RNA inhibitor and the interacting tile strand with a fluorophore and quencher pair (1, Cy3/BHQ1 and 2, Cy5/BHQ2).*

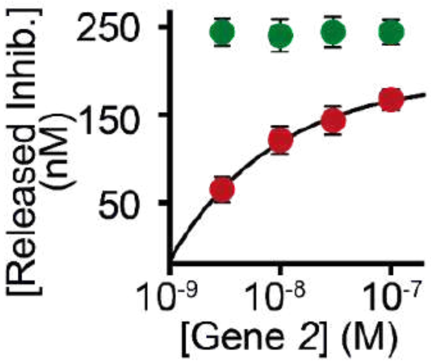


***Supplementary Figure 18.*** *Release of RNA inhibitors 1 (green) and 2 (red) from the corresponding DNA tiles after 300 min as a function of gene 2 concentration.*

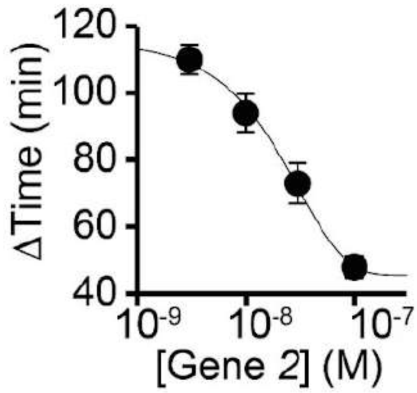


***Supplementary Figure 19.*** *Delay time (ΔT) of tile 2 activation as a function of high yield gene 2 concentration.*

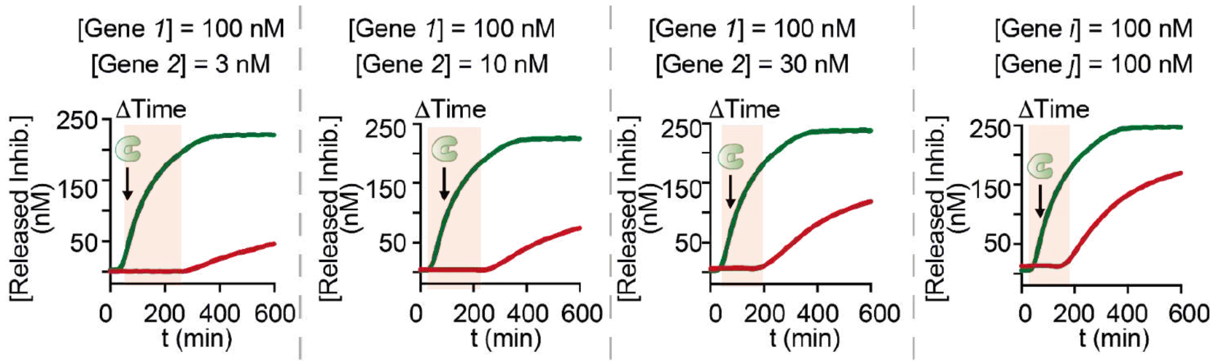


***Supplementary Figure 20.*** *Kinetic traces showing the release of RNA inhibitors (1, green and 2, red) from the corresponding tiles at different concentrations of low yield gene 2 (from 3 to 100 nM), fixed concentration of gene 1 (100 nM), connector complex (300 nM), and T7 RNAP (4 U/μL). To easily follow the release of the inhibitors after the production of the RNA activator, we modified the RNA inhibitor and the interacting tile strand with a fluorophore and quencher pair (1, Cy3/BHQ1 and 2, Cy5/BHQ2).*

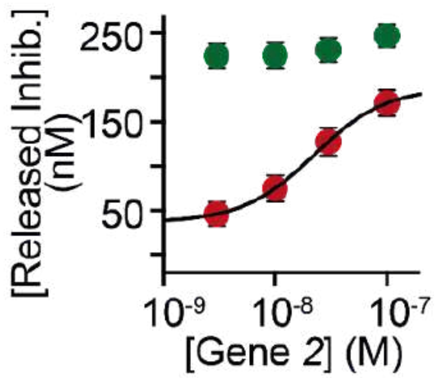


***Supplementary Figure 21.*** *Release of RNA inhibitors 1 (green) and 2 (red) from the corresponding DNA tiles after 600 min as a function of low yield gene 2 concentration.*

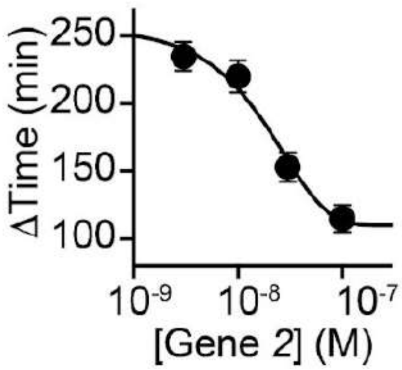


***Supplementary Figure 22.*** *Delay time (ΔT) of tile 2 activation as a function of low yield gene 2 concentration.*

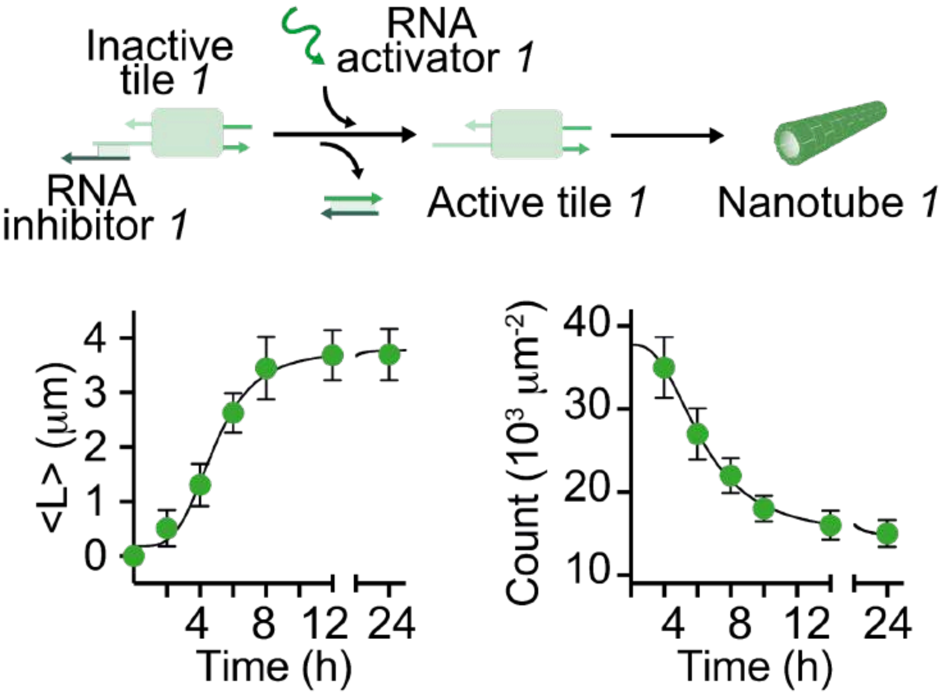


***Supplementary Figure 23. Top.*** *Schematic representation of a control experiment in which the inactivation and activation of the tiles and thus the growth of the nanotubes is mediated by the presence of synthetic RNA strands (i.e. inhibitor or activator).*

***Bottom left.*** *Kinetic traces of nanotube length measured from fluorescence microscopy images.<L>indicates the mean nanotube length of nanotubes.* **Bottom right.** *Count (number of structures per 100 μm²) of assembled green nanotubes measured from fluorescence microscopy images. Experiments shown in this Fig. were performed in 1X TXN buffer (5X contains: 200 mM Tris-HCl, 30 mM $MgCl_2$, 50 mM DTT, 50 mM NaCl, and 10 mM spermidine), pH 8.0, 30 °C. [Tile G, 1] = 250 nM; [RNA inhibitor 1] = 1 μM; [RNA activator 1] = 3 μM.*

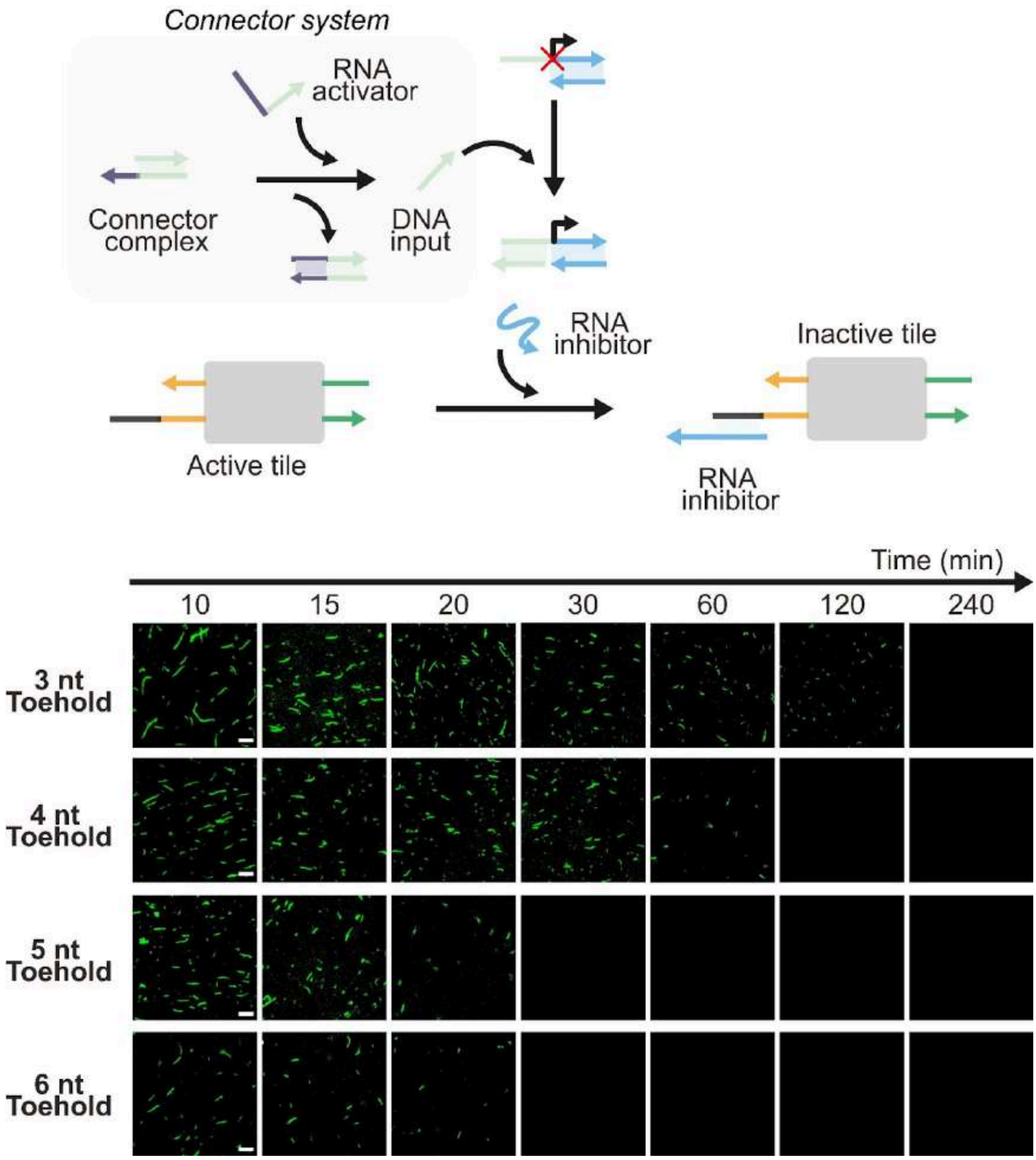


***Supplementary Figure 24. Top.*** *Schematic representation of tile inactivation mediated by the presence of a transcribed RNA inhibitor.* ***Bottom.*** *Kinetics of disassembly of tile G can be modulated by the length of the toehold of one of the strands forming the connector complex. Experiments were performed in the presence of active tiles G (250 nM), connector complex (300 nM), RNA activator (1 μM), inhibitor gene (100 nM). Fluorescence images scale bar, 2.5 μm*

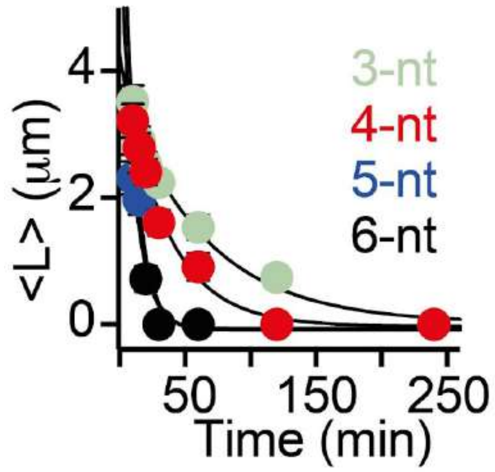


***Supplementary Figure 25.*** *Nanotube mean length at different lengths of the toehold portion (3-nt, green; 4-nt, red; 5-nt, blue; 6-nt, black) of one of the strands of the connector complex vs time (see SI 23). <L> indicates the mean nanotube length of nanotubes.*

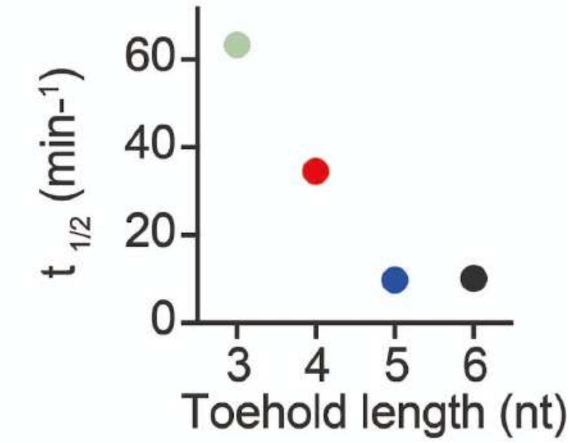


**Supplementary Figure 26.** Reaction half-life ($t_{1/2}$) of tile inhibition as a function of toehold length (from 3 to 6-nt).

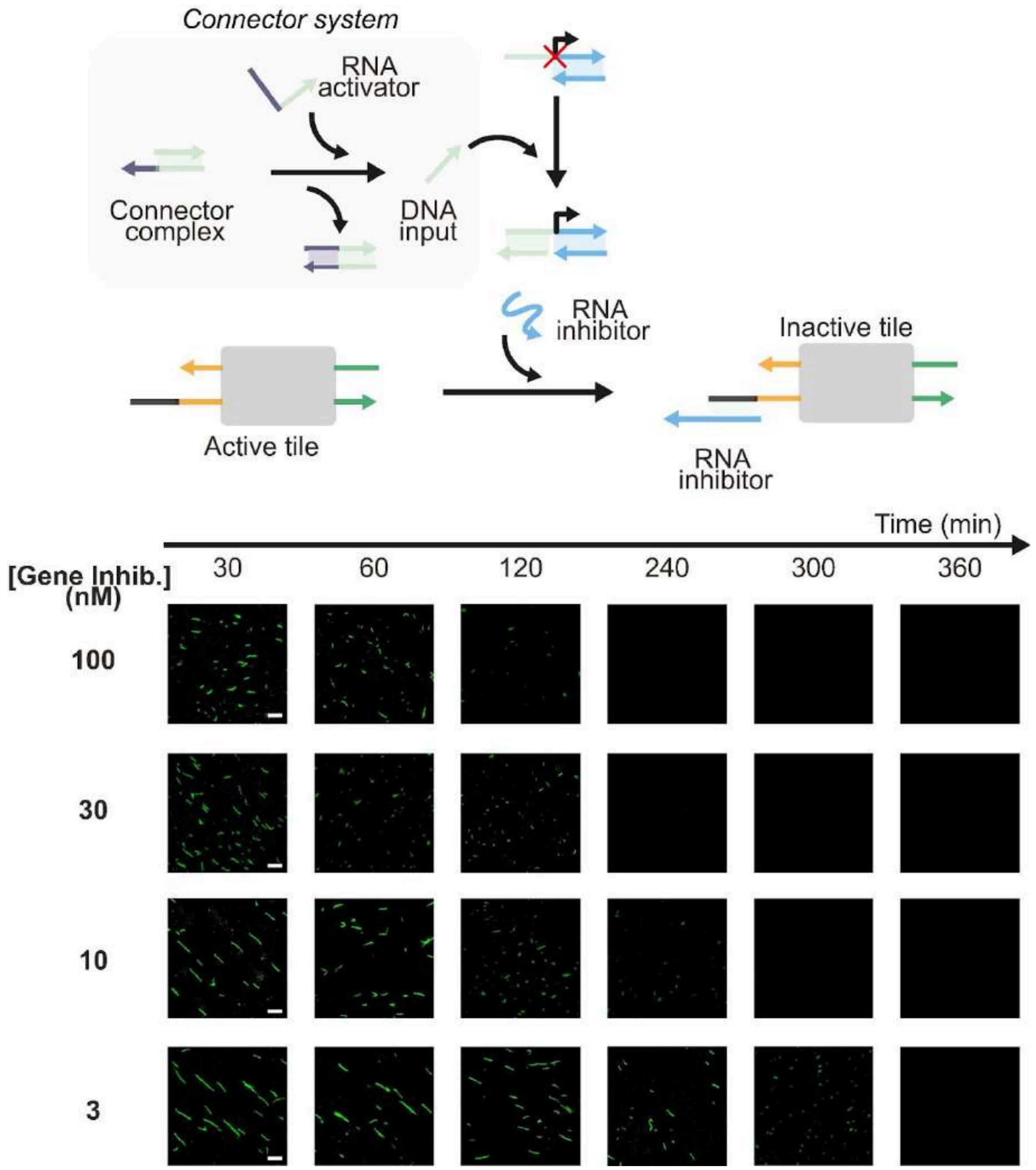


***Supplementary Figure 27. Top.*** *Schematic representation of tile inactivation by the presence of a transcribed RNA inhibitor.* ***Bottom.*** *The kinetics of degradation of the active G tiles can be modulated by the concentration of the gene producing the RNA inhibitor by using 3-nt toehold of one of the strands forming the connector complex. Experiments were performed in the presence of active tiles G (250 nM), connector complex (300 nM), RNA activator (1 µM), inhibitor gene (3-100 nM). Fluorescence images scale bar, 2.5 µm.*

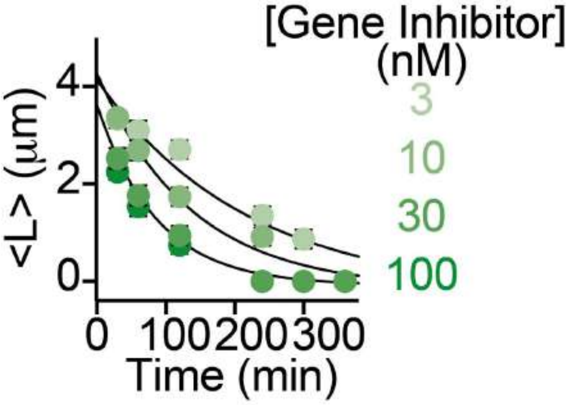


***Supplementary Figure 28.*** *Kinetic traces of nanotubes mean length. The rate of tile disassembly depends on the concentration of the gene transcribing the RNA inhibitor strand (from 3 to 100 nM; light to dark green). <L> indicates the mean nanotube length of nanotubes.*

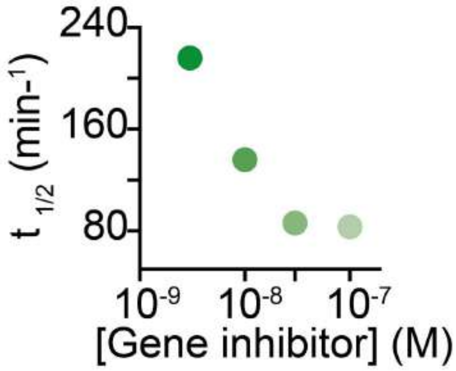


***Supplementary Figure 29.*** *Reaction half-life ($t_{1/2}$) of tile inhibition as a function of gene inhibitor concentration.*

#### Supplementary Note 1: Modeling

We modeled tile inhibition and activation through the following equivalent chemical reactions. Here species $T$ represents active tiles, $T^*$ represents inactive tiles, $I$ is the RNA inhibitor, and $R$ is the RNA activator:

$$T + I \xrightarrow{k_{TI}} T^*$$
$$T^* + R \xrightarrow{k_{TR}} T + IR$$
$$I + R \xrightarrow{k_{IR}} IR$$

We assume conservation of mass: $T^{tot} = T + T^*$, $I^{tot} = I + T^* + RI$.

To model *in situ* production of activator $R$ through *in vitro* transcription through a gene $G$ and RNA polymerase $RNAP$, we used the following reactions:

$$G + RNAP \xrightarrow{k^+} GRp \xrightarrow{k^-} G + RNAP$$
$$GRp \xrightarrow{k_{cat}} G + RNAP + R$$

When simulating two interconnected genes and tile systems (Fig. 2 of the manuscript), we used the reactions above, plus reactions modeling the release of DNA activators $A_1$ from connector complex $S_1$, and activation of gene $G_2$ through the following reactions:

$$R_1 + S_1 \xrightarrow{k_{RS}} A_2 + S_1^*$$
$$A_2 + G_2^* \xrightarrow{k_{AG}} G_2$$

where $S_1^*$ and $G_2*$ are respectively released connector complex and inactive gene. We assume mass conservation $S_1^{tot} = S_1 + S_1^*$ and $A_2^{tot} = S_1^{tot} = S_1 + A_2 + G_2$
The two independent sets of tiles, inhibitors, activators, connectors and genes were modeled through distinct species $T_1$, $I_1$, $R_1$, $S_1$, $G_1$ and $T_2$, $I_2$, $R_2$, $G_2$, assuming mass conservation for tiles, inhibitors, and genes. Tile inhibition/activation reactions were assumed to be the same for each subsystem, and rate constants were also assumed to be the consistent.

We used the reactions above to generate systems of ODEs, which were numerically integrated using MATLAB. Simulations including *in vitro* transcription assume zero initial amount of activator RNA. Other initial conditions are consistent with the experimental settings captured by the simulation. Total concentrations of templates, inhibitors, and genes were set to be consistent with experiments. We compared simulations with tile 1 activation experiments only (as tile 2 experiments yield similar results), reported in Supplementary Fig.s 30, 31, and 32. The simulations of the interconnected genes and tiles were used to generate the estimates for half-time and delay reported in Fig. 3 of the manuscript.

Reaction rate parameters were estimated from values provided in the literature[7–10]: $k_{TR} = 5 \cdot 10^5/M/s$, $k_{TI} = 1 \cdot 10^2/M/s$, $k_{IR} = 3 \cdot 10^2/M/s$, $k_{RS} = 4 \cdot 10^4/M/s$, and $k_{AG} = 3 \cdot 10^3/M/s$. Low values for the hybridization of free $T$ and $I$, and free $R$ and $I$ are based on the experimental control experiments in Supplementary Fig. 4, reported also in Supplementary Fig. 30 for comparison with the model. To reproduce experiments in Fig. 3d and e of the manuscript, in which the toehold length of the connector complex varies, we used the release rate constant $k_{RS}$ reported above to simulate the 6 nt toehold; for shorter toeholds, we scaled this constant by a factor $0.7$, $0.03$, $0.002$ respectively for 5, 4, and 3 nt toeholds.

For in vitro transcription, we used the following parameters when simulating the high yield genes: $k^+ = 1 \cdot 10^6/M/s$, $k^- = 1 \cdot 10^{-3}/s$, and $k_{cat} = 1 \cdot 10^{-3}/s$. For the low yield genes, we assumed $k^+ = 1 \cdot 10^5/M/s$ (slower binding when compared to the high yield case), $k^- = 1 \cdot 10^{-3}/s$ (faster unbinding), and $k_{cat} = 1 \cdot 10^{-3}/s$. As experiments in Fig. 2 of the manuscript show that 3 nM of gene are sufficient to release more than 30% of the inhibitor, and that addition of large amount of gene does not majorly change the *speed* of activation, we infer that the Michaelis-Menten constant of the enzyme $K_M = (k^- + k_{cat})/k^+$ is low, hence our choice of large $k^+$ when compared with the literature[9,10]. From preliminary simulations, we estimated the concentration of $RNAP$ to be $150\, nM$ when supplying 4 units of enzyme, which is not too far from previous estimates[9,10]. This concentration was scaled linearly when less units were added in Fig. 2 of the manuscript. However, the experiments reported in Supplementary Figure 32 show slow kinetics of activation when compared to the gene variation experiments in Supplementary Figure 31; these results could be explained only by assuming a less concentrated and less efficient batch of enzymes, thus we assumed that 4 units correspond to $75\, nM$ and $k^+ = 0.5 \cdot 10^6/M/s$. Where multiple genes are present, we assume the enzyme binding/unbinding and catalytic activity are the same, and that the genes compete for enzyme binding.

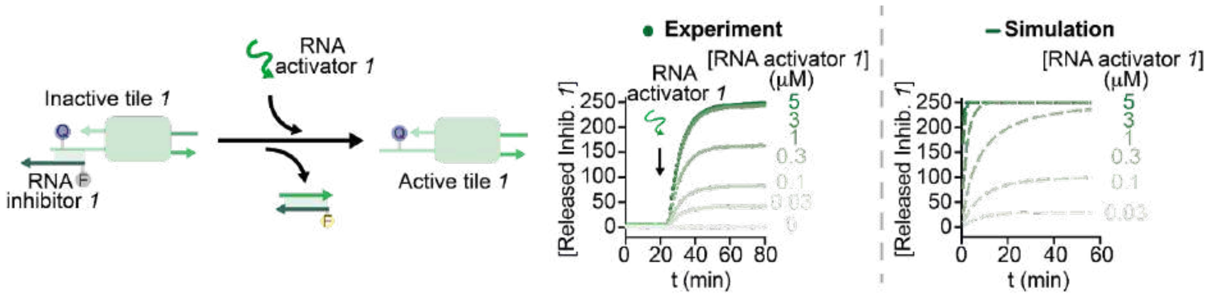


***Supplementary Figure 30.*** *Tile 1 activation using synthetic RNA activator 1 supplied manually. Left: schematic of the reaction. Right: Experimental results and simulations with reaction rates listed in the text. Experiments were performed at 30° C in 1X transcription buffer, 10 mM NTPs, pH 8.0 in a 100 µL cuvette. Experimental values are averages of three separate measurements and error bars reflect standard deviations.*

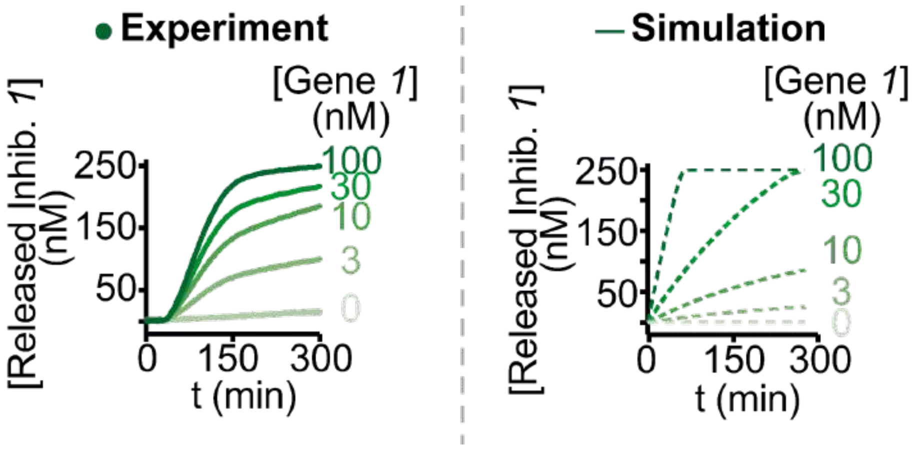


***Supplementary Figure 31.*** *Comparison of experiments and simulation results for inhibitor release when changing the amount of gene (Fig. 2c of the manuscript). Experiments were performed at 30° C, in 1X transcription buffer, 10 mM NTPs, pH 8.0 in a 100 μL cuvette. Experimental values are averages of three separate measurements and error bars reflect standard deviations.*

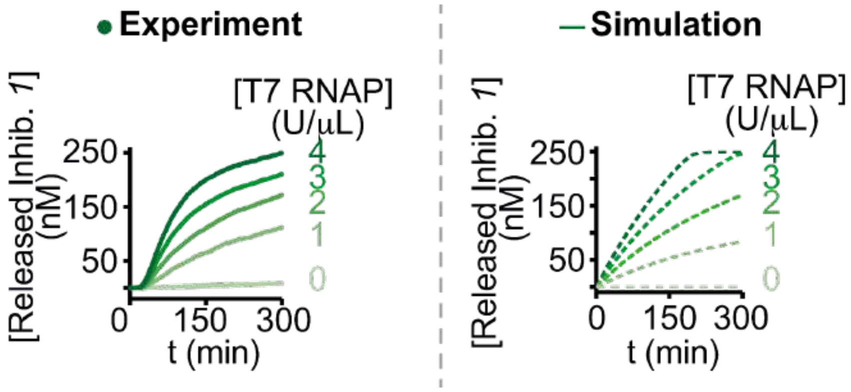


***Supplementary Figure 32.*** *Comparison of experiments and simulation results for inhibitor release when changing the amount of T7 RNAP supplied to the system (Fig. 2e of the manuscript). Experiments were performed at 30° C, in 1X transcription buffer, 10 mM NTPs, pH 8.0 in a 100 μL cuvette. Experimental values are averages of three separate measurements and error bars reflect standard deviations.*
